# A parabrachial-to-amygdala circuit that determines hemispheric lateralization of somatosensory processing

**DOI:** 10.1101/2022.09.06.506763

**Authors:** Heather N. Allen, Sarah Chaudhry, Veronica M. Hong, Lakeisha A. Lewter, Ghanshyam P. Sinha, Yarimar Carrasquillo, Bradley K. Taylor, Benedict J. Kolber

## Abstract

**Background:** The central amygdala (CeA) is a bilateral hub of pain and emotional processing with well-established functional lateralization. We reported that optogenetic manipulation of neural activity in the left and right CeA has opposing effects on bladder pain.

**Methods:** To determine the influence of calcitonin gene-related peptide (CGRP) signaling from the parabrachial nucleus (PBN) on this diametrically opposed lateralization, we administered CGRP and evaluated the activity of CeA neurons in acute brain slices as well as the behavioral signs of bladder pain in the mouse.

**Results:** We found that CGRP increased firing in both the right and left CeA neurons. Furthermore, we found that CGRP administration in the right CeA increased behavioral signs of bladder pain and decreased bladder pain-like behavior when administered in the left CeA.

**Conclusions:** These studies reveal a parabrachial-to-amygdala circuit driven by opposing actions of CGRP that determines hemispheric lateralization of visceral pain.

## Introduction

Paul Broca and Carl Wernicke introduced the phenomenon of brain lateralization in the mid 1800s by revealing that speech and language centers are predominantly located in the left cerebral hemisphere (1,2). Asymmetrical supraspinal processing is more common than previously believed and is not unique to humans (3). Brain lateralization is conserved across species and facilitates sensory, cognitive, and motor processing (3). Human neuroimaging studies reveal that brain lateralization is disturbed in the context of neurological disorders, including schizophrenia (4,5), anxiety (6,7), depression (8–10), post-traumatic stress disorder (11), and chronic pain (12,13).

Chronic pain affects approximately 35.5% of the world population (14). The incidence of affective comorbidities that appear along with the presentation of chronic pain (15) suggests the involvement of the central nervous system in the modulation of the disease. Neuroimaging studies implicate many supraspinal sites in chronic pain involvement, including the central amygdala (CeA) (16,17). The CeA is a hub of both pain and emotional processing (18), and while the lateralized functions of the CeA in the context of emotion has been known for decades (19), only recently has amygdala lateralization in the context of pain been reported (13,20,21). Hemispheric left-right differences are found in the amygdala in the context of pain in humans (17,22,23) and rodents (20,21,24). The right CeA has long been recognized as the predominate modulator of pain compared to the left CeA (20,21,25–27). The right amygdala is the major pro-nociceptive modulator in neuropathic (28), inflammatory (20,25,26), and arthritis pain (21), while the left amygdala contributes to pain modulation less or only in certain circumstances (29). However, much of what we know about amygdala lateralization in the context of pain modulation in rodents comes from studies where the injury or stimulus is restricted to a single side of the body, making it difficult to interpret the results in the context of lateralization due to the decussation of spinal neurons and the predominance of the contralateral brain to single-sided peripheral injury (30). To circumvent these limitations, we probed hemispheric lateralization using a mouse model of bladder pain.

Clinically, bladder pain conditions are categorized under the umbrella of urologic chronic pelvic pain syndrome (UCPPS). UCPPS is debilitating and predominantly affects women (31–33). Although the bladder is a bilaterally innervated midline organ (34), the left and right CeA do not contribute equally to the modulation of bladder pain in humans (34–36). Rodent studies also expose functional differences in the contribution of the left and right CeA to the modulation of bladder pain, with the right CeA serving a pro-nociceptive function and the left CeA serving an anti-nociceptive function (24). The mechanisms surrounding the existence of these left-right differences are poorly understood. Pro-nociceptive outputs from the right amygdala are associated with multiple molecular mediators (24,26,27,38–40), but the molecular mediators of left amygdala anti-nociception in lateralization remain a mystery. In numerous cases, single neuropeptides or receptors that drive right amygdala pro-nociception (e.g. metabotropic glutamate receptor 5 (mGluR5), pituitary adenylyl cyclase activating peptide (PACAP), dynorphin) have no effect in the left amygdala despite the presence of receptors and/or activity dependent changes in peptide expression (24,26,38).

The CeA receives nociceptive input from the periphery via the parabrachial nucleus (PBN) along the spino-parabrachio-amygdaloid pathway (41). The PBN is a key node in the modulation of pain and aversion (42,43). Parabrachial neurons that project to the CeA express high levels of calcitonin gene-related peptide (CGRP) (44). These neurons are implicated in visceral malaise, aversion, appetite, threat, and pain (45). Here, we investigated the contribution of parabrachial CGRP signaling in the left and right CeA to the lateralized modulation of bladder pain utilizing cell-type specific excitatory and inhibitory optogenetics and pharmacology in a mouse model of bladder pain.

## Methods

Additional details for methods can be found in *Supplemental Methods* section.

### Animals

Experiments used *Calca*^*tm1*.*1(cre/EGFP)Rpa*^ (*Calca*^*Cre*^) Cre-recombinase knockin mice, *Calcrl*^Cre^ crossed with a *Rosa26-flox-stop-tdTomato* reporter line Ai9 to generate *Calcrl*^Cre^::Ai9, or wild-type C57BL/6J female littermates. Cyclophosphamide (CYP) was used to induce a bladder pain-like sensitivity phenotype in rodents (46,47) by treating animals with CYP five days prior to experimentation.

### Stereotaxic surgeries

Adeno-associated viruses containing Cre-dependent optogenetic constructs were used to manipulate activity of *Calca*-expressing fibers from the parabrachial nucleus (PBN) in the central amygdala (CeA). Mice received a cannula or wireless LED Neurolux device (Neurolux, St. Louis, MO) in the CeA ipsilateral to PBN viral injection. In control placement experiments, cannulae were implanted over the striatum.

### Urinary bladder distention

Urinary bladder distention (UBD) and visceromotor responses (VMR) were recorded by measuring electromyography (EMG) of the external abdominal oblique muscle during noxious distention. UBD-VMR was performed as previously described (48) one to two days following the final injection of CYP under partial isoflurane anesthesia.

### Optogenetics

Light was delivered using a low-power laser diode or LED during the “light-on” timepoint. Immediately after the completion of the “light-on” timepoint, the light source was turned off and the post light timepoint was collected.

### Pharmacology

Animals received injection (1 μL) of aCSF, 100 nM CGRP, 100 nM CGRP(8-37), or a cocktail of 100 nM CGRP+100 nM CGRP(8-37) via cannula. For combined optogenetic and pharmacology experiments, animals received 1 μL of aCSF, 100 nM CGRP(8-37), or a cocktail of 22 mM AP5 and 38 mM NBQX (49) before receiving light stimulation. For knockout experiments, optogenetic and pharmacology experiments were performed in the same animals in a randomized order and no effect of order was found (**Supplementary Fig. 7**).

### Mechanical sensitivity

*In vivo* behavioral testing was conducted one to two days following final CYP injection (day 6-7). This correlates to day 20 in experiments where animals received optogenetic stimulation of CGRP-containing PBN fibers. Calibrated von Frey filaments were used to assess abdominal sensitivity on the right and left abdomen approximately 0.5 cm from the urethra via the up-down method to calculate 50% withdrawal thresholds (50).

### Real time place preference

Animals were habituated to a three-chamber place preference apparatus with distinct visual patterns. The next day (day 20 post-surgery), animals were then placed back in the place preference apparatus where one chamber was tuned for wireless Neurolux LED stimulation. Upon entering the tuned chamber, the Neurolux device automatically started stimulation, which ended as soon as the animal exited the Neurolux tuned chamber. Animals’ activity was video recorded for 20 min using AnyMaze.

### Immunohistochemistry

All viral constructs used in these experiments contained an mCherry sequence to allow for viral targeting in *Calca-*expressing cells in the PBN and terminals in the CeA. To quantify CGRP, brains were processed for CGRP immunohistochemistry in representative sections from across the rostral-caudal axis of the CeA. All microscope images were acquired using settings from a negative control and settings were kept consistent. Fluorescence intensity of the 488 channel was normalized to fluorescence intensity of the DAPI channel for each image. The CeC was defined as the area 200 μm inward from BLA/CeA border (24).

### Electrophysiology

Whole-cell current clamp recordings were restricted to late-firing fluorescently labeled neurons expressing the CGRP receptor (CGRPR+) in slices from *Calcrl*^Cre^::Ai9 mice or unlabeled neurons in slices from C57BL/6J wild-type mice within the capsular subdivision of the CeA.

### RNAscope *in situ* hybridization

RNAscope probes for *Calcrl, Prkcd*, and *Sst* were used in representative sections from across the rostral-caudal axis of the CeA. Positive cells were identified as a DAPI-labeled nucleus surrounded by at least three puncta. Cell counts were determined blinded to treatment. Cell number and percent co-localization were averaged across sections from the same brain.

### CeA tissue collection and cAMP ELISA

Wild-type C57BL/6J mice received CGRP or aCSF infused into the bilateral CeA via cannula. Mice were decapitated 40 minutes later, and brains were sectioned, flash frozen, and homogenized prior to completing cAMP ELISA according to kit instructions.

### Statistics and data analysis

All data analyses were conducted blind to treatment/virus/genotype. UBD data was analyzed via unpaired *t*-tests, repeated-measures two-way analysis of variance (ANOVA) followed by Bonferroni or Dunnett’s post hoc tests. Behavioral data was analyzed using paired *t*-tests, one-way ANOVA, or repeated measures two-way ANOVAs followed by Bonferroni or Dunnett’s post hoc tests. Electrophysiology data were analyzed via two-way repeated measures ANOVA. RNAscope data was analyzed using two-way ANOVAs followed by Tukey post-hoc test. Statistical significance was determined at the level of P<0.05. Asterisks denoting P values include: *P<0.05, **P<0.01, ***P<0.001, and ****P<0.0001. All data are presented as the mean +/- standard error of the mean (SEM). Statistical information for all figures is provided in **Supplementary Table 1**.

## Results

### Left and right PBN→CeA CGRP fibers have opposing roles on bladder pain-like physiology

To explore functional lateralization of the CeA in the context of bladder pain, we optogenetically manipulated CGRP-containing PBN projecting fibers with a Cre-dependent channelrhodopsin (ChR2) expressed in CGRP-containing PBN fibers in the left and right CeA during noxious bladder distention in a cyclophosphamide (CYP)-induced mouse model of bladder pain (**Fig. 1A-C**). Visceromotor responses (VMRs) to noxious bladder distention increased in mice with CYP-induced cystitis (**Fig. 1D**). Optogenetic activation of CGRP-containing PBN fibers in the left CeA decreased VMRs to noxious UBD, suggesting that activation of these terminals reduced bladder pain-like physiology in CYP-treated mice (**Fig. 1E-G**). In contrast, optogenetic activation of right PBN→CeA CGRP terminals further increased VMRs (**Fig. 1H-J**).

**Figure 1:**
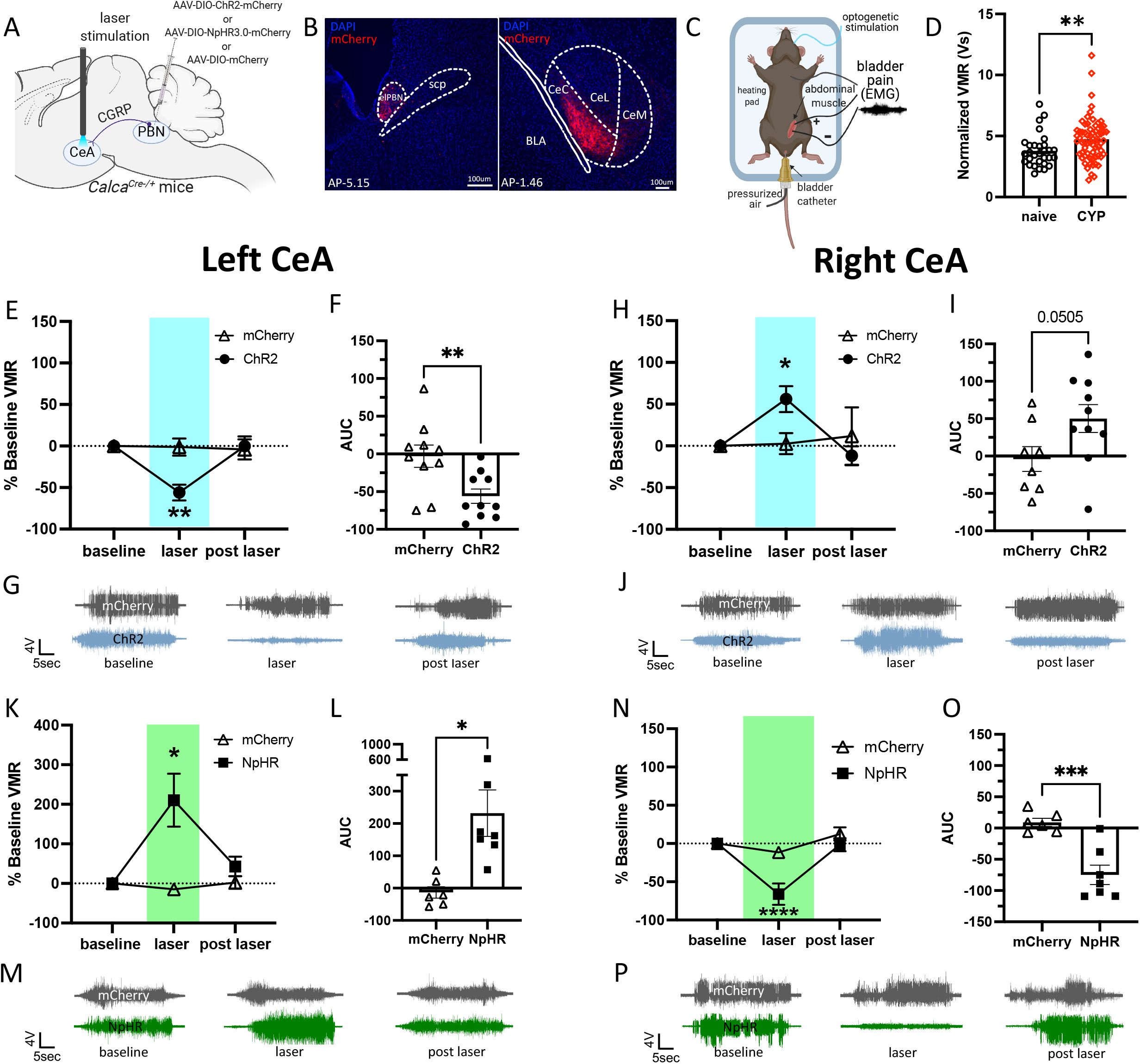
Optogenetic stimulation of parabrachial CGRP terminals in the left and right CeA has opposing effects on bladder pain-like physiology. **A**) Schematic of surgical set up for optogenetic activation or inhibition of CGRP positive PBN terminals in the left or right CeA. **B**) Representative images of mCherry labeling CGRP positive cell bodies in the PBN (left) and terminals in the CeA (right). **C**) Schematic for UBD-VMR recording during optogenetic stimulation of CGRP fibers in the left or right CeA. **D**) Normalized VMRs to 60 mmHg distention in naïve and CYP-treated mice. **E**) Percent change from baseline VMRs to 60 mmHg distention during and after optogenetic activation of CGRP terminals in the left CeA. **F**) Area under the curve (AUC) for (**E**). **G**) Representative EMG traces for baseline, light on, light off timepoints in (**E**). **H**) Percent change from baseline VMRs to 60 mmHg distention during and after optogenetic activation of CGRP terminals in the right CeA. **I**) AUC for (**H**). **J**) Representative EMG traces for baseline, light on, light off timepoints in (**H**). **K**) Percent change from baseline VMRs to 60 mmHg distention during and after optogenetic inhibition of CGRP terminals in the left CeA. **L**) AUC for (**K**). **M**) Representative EMG traces for baseline, light on, light off timepoints in (**K**). **N**) Percent change from baseline VMRs to 60 mmHg distention during and after optogenetic inhibition of CGRP terminals in the right CeA. **O**) AUC for (**N**). **P**) Representative EMG traces for baseline, light on, and light off timepoints for (**N**). All data are presented as mean +/- SEM and error bars represent SEM. *P<0.05, **P<0.01, ***P<0.001. See supplementary table 1 for further statistical information.

Halorhodopsin (NpHR)-mediated optogenetic inhibition of CGRP-containing PBN→CeA terminals had opposing effects in the left and right CeA. Silencing CGRP terminals in the left CeA increased VMRs, (**Fig. 1K-M**), while optogenetic inhibition of CGRP terminals in the right CeA decreased VMRs in CYP-treated mice (**Fig. 1N-P**). These experiments demonstrate that left versus right PBN→CeA CGRP-expressing terminals have opposing effects on the modulation of bladder pain.

### Activation of PBN→CeA CGRP-containing terminals mediates bladder sensitivity but not affective pain

We utilized Cre-dependent ChR2-mCherry (or control mCherry only) expression in the left or right PBN of *Calca*-Cre mice and wireless blue LED (Neurolux) devices to investigate the effect of activating PBN→CeA CGRP terminals in the left and right CeA on pain-like behaviors in awake, freely moving animals (**Fig. 2A-C**). Optogenetic activation of CGRP terminals in the left CeA increased 50% withdrawal thresholds while activation of CGRP terminals in the right CeA decreased 50% withdrawal thresholds (**Fig. 2D, F**). CYP increases abdominal mechanical sensitivity so severely (**Fig. 2C**) that changes in 50% withdrawal thresholds are difficult to observe post CYP induction. For this reason, 50% withdrawal thresholds were also analyzed as a percent baseline to more clearly demonstrate the optogenetic-induced changes in abdominal sensitivity (**Fig. 2E, G**). These findings recapitulate the results observed in lightly anesthetized animals during UBD (**Fig. 1**), further demonstrating that CGRP-expressing terminals in the left and right CeA differentially modulate bladder pain-like behavior.

**Figure 2:**
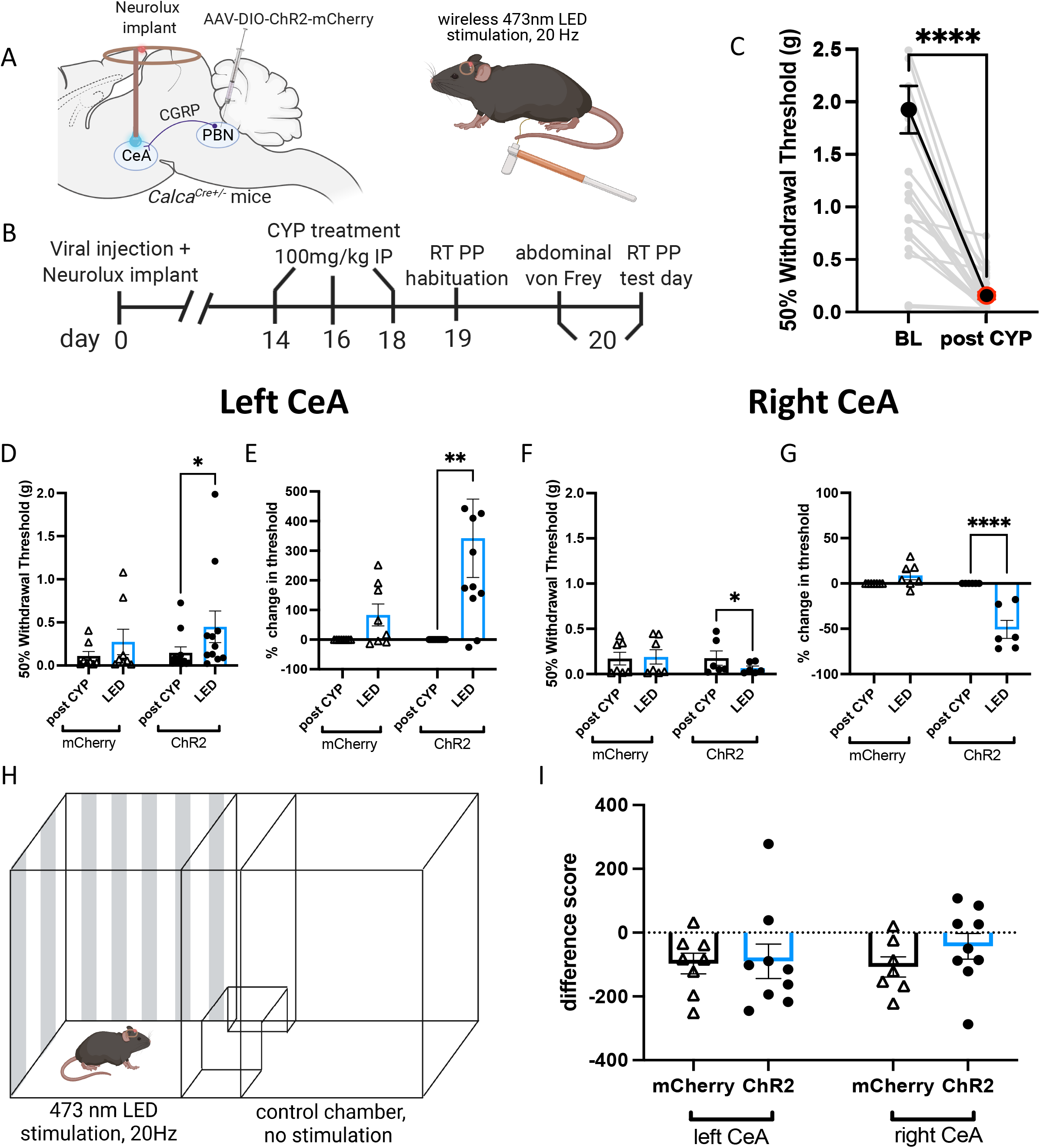
Optogenetic activation of parabrachial CGRP terminals in the left and right CeA has opposing effects on pain-like behavior. **A**) Schematic for wireless optogenetic activation of PBN CGRP terminals in the left or right CeA during abdominal von Frey. **B**) Timeline for wireless optogenetic activation of CGRP terminals during behavior in awake animals. **C**) Abdominal 50% withdrawal thresholds before and after CYP treatment. **D**) 50% withdrawal thresholds before and during optogenetic activation of CGRP terminals in the left CeA of CYP mice. **E**) Percent change in 50% withdrawal threshold from (**D**). **F**) 50% withdrawal thresholds before and during optogenetic activation of CGRP terminals in the right CeA of CYP mice. **G**) Percent change in 50% withdrawal threshold from (**F**). **H**) Schematic for wireless optogenetic activation of CGRP terminals in the left or right CeA during real time place preference/aversion. **I**) Difference scores for CYP animals with ChR2 or mCherry during real time place preference/aversion. All data are presented as mean +/- SEM and error bars represent SEM. *P<0.05, **P<0.01, ****P<0.0001. See supplementary table 1 for further statistical information.

To investigate the influence of PBN→CeA CGRP-expressing terminals on pain-related aversion, we evaluated real-time place preference/aversion using the same CYP-treated animals containing Cre-dependent ChR2 in the left or right PBN and ipsilateral CeA Neurolux implants. Mice were stimulated in one of two chambers during a 20 min trial (**Fig. 2H**) and showed no preference or aversion to the LED-associated chamber (**Fig. 2I**). Our results reveal the PBN projecting CGRP-containing terminals in the left and right CeA differentially mediate bladder pain-like sensation but not pain-related acute aversion.

### CGRP increases the firing rate of neurons in the right and left CeA

To assess the effect of CGRP on the activity of CeA neurons, we performed whole-cell patch-clamp recordings from left and right CeA neurons in acute brain slices obtained from naïve mice (**Fig. 3A**). The number of action potentials elicited by depolarizing current injections of various amplitudes was measured after bath application of CGRP (500 nM) or aCSF. In all recordings, the number of action potentials increased as a function of the amplitude of depolarizing current injected (**Fig. 3C-H**). Action potential firing and resting membrane potential in CeA neurons recorded in the both the left and right hemisphere were significantly higher after CGRP as compared to aCSF (**Fig. 3C-J**), demonstrating that CGRP-mediated increases in action potential firing are not lateralized in the CeA.

**Figure 3:**
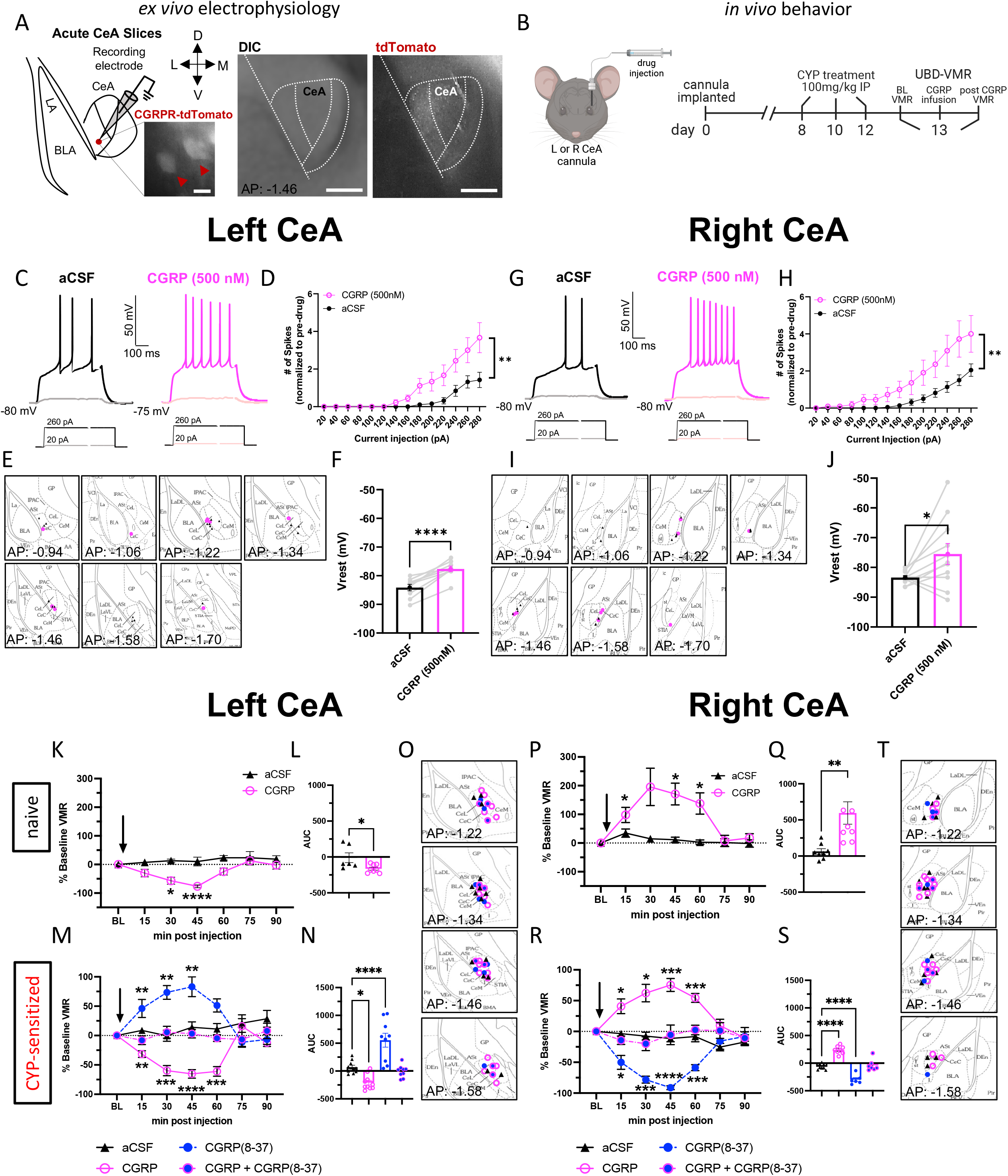
CGRP pharmacology in the left and right CeA. **A**) Schematic and representative image of recording site in the CeA. Scale bars represent 200 μm (left image) and 250 μm (right imags) **B**) Schematic and timeline for CGRP pharmacology in the left or right CeA during UBD-VMR. **C**) Representative traces of firing patterns of left CeA neurons after aCSF or CGRP perfusion. **D**) Number of spikes of left CeA neurons after aCSF or CGRP perfusion, normalized to before bath exchange. **E**) Targeting of left CeA neurons recorded. **F**) V_rest_ of left CeA neurons after aCSF or CGRP perfusion. **G**) Representative traces of firing patterns of right CeA neurons after aCSF or CGRP perfusion. **H**) Number of spikes of right CeA neurons after aCSF or CGRP perfusion. **I**) Targeting of right CeA neurons recorded. **J**) V_rest_ of right CeA neurons after aCSF or CGRP perfusion. **K**) Percent change from baseline VMRs to 60 mmHg distention after infusion of aCSF or CGRP into the left CeA of naive mice. **L**) AUC for (**K**). **M**) Percent change from baseline VMRs to 60 mmHg distention after infusion of aCSF, CGRP, CGRP(8-37), or CGRP+CGRP(8-37) into the left CeA of CYP mice. **N**) AUC for (**M**). **O**) Targeting of cannulas in the left CeA. **P**) Percent change from baseline VMRs to 60 mmHg distention after infusion of aCSF or CGRP into the right CeA of naive mice. **Q**) AUC for (**P**). **R**) Percent change from baseline VMRs to 60 mmHg distention after infusion of aCSF, CGRP, CGRP(8-37), or CGRP+CGRP(8-37) into the right CeA of CYP mice. **S**) AUC for (**R**). **T**) Targeting of cannulas in the right CeA. All data are presented as mean +/- SEM and error bars represent SEM. *P<0.05, **P<0.01, ****P<0.0001. See supplementary table 1 for further statistical information.

### Pharmacological activation of CGRP receptors in the left and right CeA differentially affects bladder pain-like physiology

The robust and distinctive effects that CGRP-expressing terminals in the left versus right CeA have on bladder pain prompted us to investigate whether CGRP itself influences CeA lateralization. We utilized CGRP pharmacology in the left or right CeA to record VMRs of wild-type naïve or bladder-sensitized mice following intra-CeA injection of artificial cerebrospinal fluid (aCSF), CGRP, the peptide antagonist CGRP(8-37), or a cocktail of CGRP+CGRP(8-37). In naïve animals, CGRP decreased pain-like responses to UBD when infused in the left CeA (**Fig. 3K-L**) but increased pain-like responses when infused in the right CeA (**Fig. 3P-Q**). This pattern of lateralization was maintained in CYP-sensitized animals (**Fig. 3M-T**). VMRs did not change when equal parts CGRP and CGRP(8-37) were infused together, nor after vehicle infusion. In a placement control experiment targeting the striatum of naïve mice, CGRP had no effect on VMRs (compared to pre-treatment baseline) (**Supplemental Fig. 9**). Overall, these results suggest that CGRP contributes to CeA lateralization in the modulation of physiological responses to noxious bladder stimulation under both naïve and injured conditions.

### CGRP drives CeA optogenetic lateralization in the context of bladder pain

PBN→CeA CGRP neurons are heterogenous and express numerous neurotransmitters and peptides (43,44,51,52). To confirm that CGRP is the driving force behind the lateralized function of the PBN→CeA circuit in bladder pain, we used UBD to assess the effects of combining pharmacological blockade of CeA cells (aCSF, CGRP(8-37), or the glutamatergic transmission blockers AP5+NBQX) with optogenetic activation of CGRP-expressing PBN terminals in the CeA. VMRs were collected at baseline, at peak level of drug activation (49,53) during laser stimulation, and after the drug effects were gone and the laser was turned off. AP5+NBQX did not change the effects of optogenetic stimulation of left or right PBN→CeA CGRP terminals on bladder pain (**Fig. 4B-C**). Control animals receiving infusion of aCSF exhibited the same anti-hyperalgesic and hyperalgesic effects (**Fig 4B-C**) of optogenetic CGRP terminal activation in the left and right CeA, respectively, as observed in previous experiments (**Fig. 1E-I**). CGRP(8-37), however, blocked the effects of optogenetic stimulation on bladder pain-like physiology (**Fig. 4B-C**), suggesting that CGRP release is responsible for optogenetic-induced lateralized changes in bladder pain.

**Figure 4:**
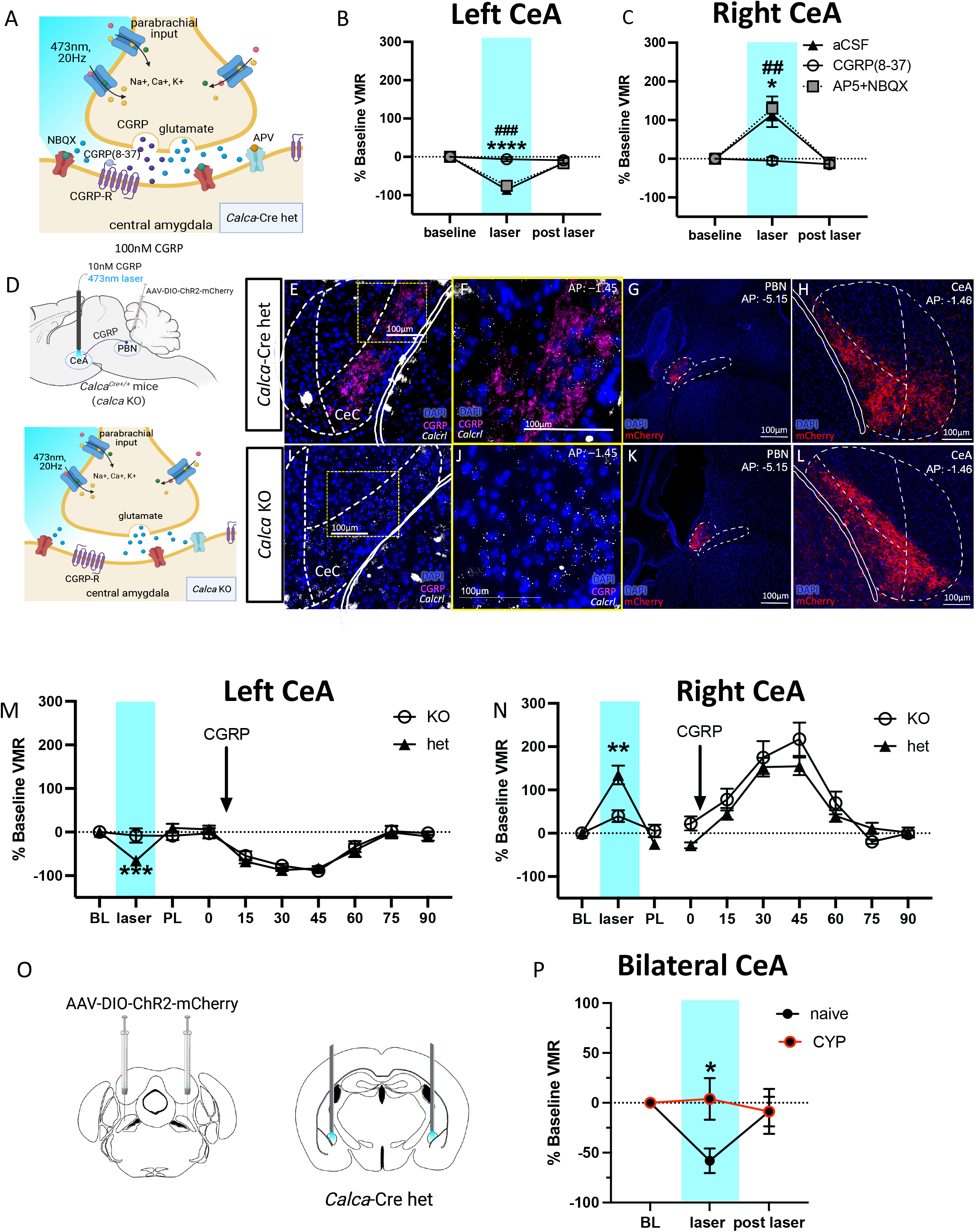
Effects of optogenetic activation in the left and right CeA is due to parabrachial CGRP signaling. **A**) Schematic for blocking CGRP receptors or AMPA and NMDA receptors in the CeA during optogenetic activation of CGRP terminals from the PBN. **B**) Percent change in VMRs to 60 mmHg distention at baseline, during optogenetic activation and pharmacological inhibition, and after optogenetic and pharmacological (# AP5+NBQX, *aCSF) manipulation in the left CeA. **C**) Percent change in VMRs to 60 mmHg distention at baseline, during optogenetic activation and pharmacological inhibition, and after optogenetic and pharmacological manipulation in the right CeA. **D**) Schematics for optogenetic activation of CGRP terminals and pharmacological activation of CGRP receptor cells in *Calca-*Cre heterozygous and knockout mice. **E, F, I, J**) Representative immunohistochemistry for CGRP (fuchsia) and RNAscope for *Calcrl* (white) in the CeA of *Calca*^Cre/+^ heterozygous (**E-F**) and *Calca*^Cre/Cre^ homozygous (**I-J**) mice. **G, H**) Representative mCherry tagged ChR2 in Cre positive cells the PBN (**G**) and terminals in the CeA (**H**) of *Calca-*Cre heterozygous mice. **K, L**) Representative mCherry tagged ChR2 in Cre positive cells in the PBN (**K**) and terminals in the CeA (**L**) of homozygous mice. **M**) Percent change from baseline VMRs to 60 mmHg distention during optogenetic activation of CGRP terminals (left) and pharmacological activation or CGRP receptor cells (right) in the left CeA of *Calca*-Cre heterozygous and homozygous CYP mice. **N**) Percent change from baseline VMRs to 60 mmHg distention during optogenetic activation of CGRP terminals (left) and pharmacological activation of CGRP receptor cells (right) in the right CeA of *Calca*^Cre^ heterozygous and homozygous CYP mice. **O**) Schematic for bilateral ChR2 injections in PBN and cannulae in CeA of *Calca*^*Cre*^ mice. **P**) Percent change from baseline VMRs to 60 mmHg pressure during bilateral optogenetic stimulation of CGRP-containing PBN→CeA terminals in naïve and CYP-treated animals. All data are presented as mean +/- SEM and error bars represent SEM. *P<0.05, **P<0.01, ***P<0.001 ****P<0.0001. See supplementary table 1 for further statistical information.

To further confirm that CGRP drives CeA lateralization, we utilized a combination of the same Cre-dependent optogenetic and pharmacological activation approaches used in the previous experiments during UBD in *Calca*^*Cre*^ heterozygous and *Calca*^*Cre/Cre*^ homozygous (“CGRP-knockout”) animals (**Fig. 4D-L**). CYP-treated animals underwent noxious UBD to record baseline VMRs, followed by optogenetic activation of ChR2-expressing PBN→CeA terminals in the left or right CeA and pharmacological stimulation via CGRP infusion into the right or left CeA. Every animal received both optogenetic and pharmacological activation administered in a randomized order; we found no significant effect of the order of activation (**Supplementary Fig. 7**). Optogenetic stimulation did not change bladder pain-like responses in CGRP-knockout animals, whereas *Calca*^Cre^ heterozygous animals showed the same anti-hyperalgesic and hyperalgesic effects (**Fig. 4M-N**) observed in earlier optogenetic experiments (**Fig. 1E-I**; **Fig. 4B-C**) in the left and right CeA, respectively. While the knockout genotype prevented optogenetic-induced changes, there was no difference between genotypes in response to CGRP infusion; both CGRP-knockout and heterozygote animals displayed a decrease in bladder pain-like physiology when CGRP was infused into the left CeA and an increase in pain-like physiology when infused into the right CeA (**Fig. 4M-N**).

### The balance of CGRP-driven CeA laterization shifts in the context of bladder injury

Finally, we evaluated how the pro and anti-nociceptive CGRP-driven functions of the left and right CeA coordinate to modulate bladder pain in naïve animals as well as how the balance changes in the context of CYP-induced bladder sensitization. We used ChR2 to bilaterally stimulate CGRP-containing PBN projecting terminals in the CeA during UBD-VMR (**Fig. 4O**). Bilateral optogenetic stimulation decreased VMRs from baseline in naïve animals but had no effect in animals with CYP-treated animals (**Fig. 4P**). These data suggest that the CGRP-mediated anti-nociceptive drive of the left CeA is stronger in naïve animals than in those with CYP-induced bladder sensitization.

To explore possible anatomical differences stemming from expression of CGRP and/or the CGRP receptor, we used immunohistochemistry to label and quantify CGRP content in the left and right CeA of saline and CYP-treated mice (**Fig. 5A-D**). We found that CYP-treated mice had lower levels of CGRP expression, and this was seen predominantly in the left CeA (**Fig. 5E**).

**Figure 5:**
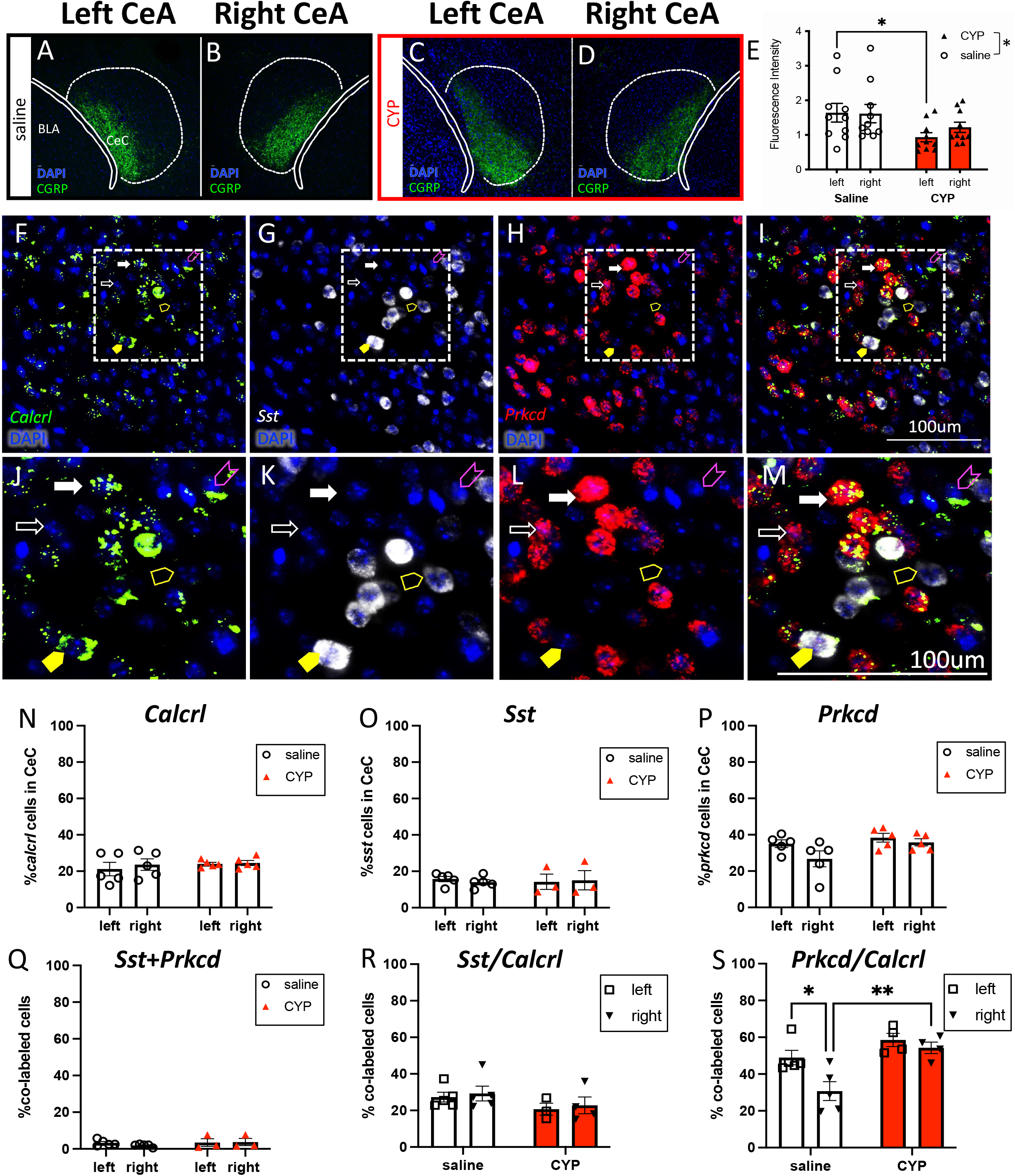
Molecular comparison of the left and right CeA in pain and non-pain mice. **A, B**) Representative immunohistochemistry for CGRP in the left (**A**) and right (**B**) CeA of saline treated mice. **C, D**) Representative immunohistochemistry for CGRP in the left (**C**) and right (**D**) CeA of CYP-treated mice. **E**) Quantified fluorescence intensity of CGRP in the left and right CeA of saline and CYP mice. **F, G, H, J, K, L**) Representative RNAscope for *Calcrl* (**F, J**), *Sst* (**G, K**), and *Prkcd* (**H, L**) in the CeA. **I, M**) Representative merge to show co-localization (fuchsia arrowhead: *Calcrl* only, yellow arrowhead: *Sst* only, white arrowhead: *Prkcd* only, filled yellow arrowhead: *Calcrl* and *Sst*, filled white arrowhead: *Calcrl* and *Prkcd*). **N**) Percent of *Calcrl*-positive cells in the left and right CeC of saline and CYP mice. **O**) Percent of *Sst*-positive cells in the left and right CeC of saline and CYP mice. **P**) Percent of *Prkcd*-positive cells in the left and right CeC of saline and CYP mice. **Q**) Percent of *Sst* and *Prkcd*-positive cells in the left and right CeC of saline and CYP mice. **R**) Percent of *Calcrl*-positive cells that also express *Sst* in the left and right CeC of saline and CYP mice. **S**) Percent of *Calcrl*-positive cells that also express *Prkcd* in the left and right CeC of saline and CYP mice. All data are presented as mean +/- SEM and error bars represent SEM. *P<0.05 **P<0.01. See supplementary table 1 for further statistical information.

### Molecular identity of CGRP receptor cells in the CeA

The capsular region of the CeA (CeC) receives the majority of PBN input and is defined by CGRP fiber expression. We investigated the CeA cells that receive CGRP input to assess if CGRP receptor expression differed in the left and right CeA of pain and non-pain animals. RNAScope fluorescence *in situ* hybridization (**Fig. 5F**) revealed the number of cells expressing *Calcrl* in the left versus right CeC of control and CYP animals did not differ (**Fig. 5N**), suggesting that *Calcrl* expression (1) does not differ between hemispheres, and (2) does not change in the context of CYP-induced bladder pain. Additionally, we found that CGRP infusion into the left or right CeA increases cAMP concentration compared to infusion of aCSF, suggesting that activation of the CGRP receptor does not differentially alter cAMP levels in the left and right CeA, consistent with the canonical G_αs_ Calcrl signaling cascade (**Supplementary Fig. 9**).

We next sought to study the molecular identity of CGRP receptor-expressing cells in the left and right CeC. We co-localized *Calcrl* with *Sst* (somatostatin, SOM) and *Prkcd (*protein kinase C delta, PKCδ), two non-overlapping targets of parabrachial projections with opposing roles in pain (40,54). There was no difference in the number of *Sst* or *Prkcd*-expressing cells between the left and right CeC nor between animals with and without CYP treatment (**Fig. 5O-P**), suggesting that core anatomical expression differences do not contribute to functional CeC lateralization in bladder pain. The percent of *Calcrl*-expressing CeC cells that also expressed *Sst* did not differ between sides or change in the context of pain (**Fig. 5R**). However, control animals showed less *Calcrl*-expressing cells that also expressed *Prkcd* in the right CeC, but this difference disappeared in the context of CYP-induced bladder sensitization; CYP animals showed no lateralization of *Calcrl*+*Prkcd* in the CeC (**Fig. 5S**). These data demonstrate that in animals without bladder pain, *Calcrl*-expressing cells have lower co-expression with *Prkcd* in the right CeC, but in the context of CYP-induced bladder sensitization, more *Calcrl* cells express *Prkcd*.

## Discussion

Our studies demonstrate a substantive functional lateralization in the PBN→CeA circuit in the context of bladder pain. Strikingly, this lateralization appears to be driven by the same neuropeptide, CGRP, in both hemispheres. We demonstrated that optogenetic manipulation of left versus right PBN→CeA CGRP terminals has opposing effects on bladder pain-like behaviors but does not influence pain-related aversion after bladder injury. Using CGRP-knockout animals, we established that it is indeed the action of CGRP released from the PBN that drives right CeA-optogenetically-mediated hyperalgesia and left CeA-optogenetically-mediated anti-hyperalgesia in bladder pain. Furthermore, our studies reveal that *Calcrl*-expressing cells in the CeA have distinct co-expression patterns with *Prkcd* that change in the context of CYP-induced bladder sensitization. To our knowledge, this is the first study not only to confirm that CGRP in the right CeA contributes to modulation of visceral pain, as it does with other pain models (55–58), but also to demonstrate that CGRP has an opposing and anti-hyperalgesic role in the left CeA in the context of bladder pain.

There is a wealth of conflicting evidence for the CeA being both pro- and anti-nociceptive across different pain models, manipulations, and experimental endpoints (13). Our right CeA CGRP data aligns with what is already known about CGRP’s well-established pro-nociceptive role in the right CeA in other pain models (55–59). Our discovery of a role for CGRP as an anti-nociceptive mediator in the left CeA in the context of bladder pain is supported by a single study that found that CGRP infused into the left CeA increased mechanical threshold in naïve rats (60). Combined with our data demonstrating diametrically opposed roles of CGRP in left and right CeA in the context of bladder pain, it is possible that CGRP in the CeA has lateralized effects in other pain models as well.

Simple anatomical differences, such as the proportion of cells expressing pro- or anti-nociceptive markers (40,53,61–63), could underlie or contribute to CeA lateralization. Recent work demonstrates bimodal modulation of neuropathic pain by SOM- and PKCδ*-*expressing cells in the right CeA (40). We observed changes in co-expression of *Prkcd* with *Calcrl* in the context of CYP. Though we are unable to confirm whether this increase in expression is biologically significant at this time, these findings should be explored in future studies. Techniques that will provide a more rigorous analysis of left versus right hemispheric differences in pain and non-pain states, such as Fos-mediated Targeted Recombination in Active Populations (FosTRAP), Fluorescence-Activated Cell Sorting (FACSorting), and RNA sequencing of Fos-labeled neurons in response to noxious stimulation will be extremely important to address this question.

In contrast to anatomical lateralization, there is evidence for time-dependent lateralization of neuronal activity in the CeA in response to pain (64). The coordination of the left and right CeA to modulate bladder pain may also have a time-dependent factor that alters the hemispheric dominance of neural activity after injury. Using bilateral optogenetic stimulation, we found that the balance of the CGRP-driven opposing functions of the left and right CeA shifts in the context of CYP. In naïve animals, the left CeA’s anti-hyperalgesic function is dominant in modulating bladder pain-like behavior, but this effect is lost in CYP-treated animals. The decreased CGRP expression in the left CeA following CYP may partially account for the loss of anti-hyperalgesia observed in CYP animals during UBD-VMR. Investigation of time-dependent hemispheric changes in neuronal activity will be important to explore as pain shifts to the more chronic state, which may provide more insight into the type of disruptions that occur in chronic pain.

In the context of the human condition, left-right differences have been observed in the amygdala of UCPPS patients. UCPPS studies using neuroimaging report lateralization of amygdala volume, activation, and functional connectivity differ in patients with and without bladder pain conditions. Women with bladder pain have increased left amygdala grey matter volume compared to women without bladder pain (36,65), and functional magnetic resonance imaging studies reveal that women with bladder pain conditions have increased connectivity between the left amygdala and periaqueductal grey (37). Changes to the typical amygdala asymmetry in the context of bladder pain suggest that altered brain structure and connectivity may exist in other types of chronic pain as well.

Our studies establish that the lateralized modulation of bladder pain-like behaviors by the CeA is driven via CGRP signaling from the PBN. Surprisingly, we found that optogenetic manipulation of PBN→CeA CGRP terminals in neither the left nor right CeA influenced pain-related aversion. Previous studies indicate that nonspecific bilateral optogenetic stimulation of PBN terminals in the CeA of normal mice induces both robust real time and conditioned place aversion (68). Interestingly, bilateral activation of CGRP-containing PBN projection terminals in the rostral but not caudal CeA produced place aversion (69). Our CeA targets were predominantly caudal, suggesting that signaling from CGRP positive PBN projection neurons to the caudal CeA modulates the sensory components of pain without regulating the aversive components. Alternatively, unilateral activation of CGRP-expressing PBN→CeA terminals may not be sufficient to modulate the aversive components of bladder pain. It is possible that both the left and right CeA are required for developing a pain-associated preference or aversion.

Advent of new technologies allows us to investigate complex processes like pain more completely via the labeling and manipulation of neural ensembles (68). CGRP signaling in the CeA contributes largely to the modulation of bladder pain-like behaviors, but it is only one part of a larger ensemble that encodes pain in the brain. Future investigation of how CGRP signaling in the CeA fits into broader circuits that modulate pain will reveal more extensive insight into the phenomenon of CeA lateralization and pain processing.

## Acknowledgments

We thank Dr. Sarah Woodley and Mauricio Gil-Silva for technical assistance and support and Dr. Richard Palmiter for providing original *Calca*^*Cre*^ and *Calcrl*^Cre^ mice. Additionally, thank you to Dr. Jordan McCall for his assistance and effort to complete electrophysiological recordings. We would also like to thank Olivia Babyok and Nina Gakii for technical assistance with the ELISA and Uma Chatterjee with brain processing. This work was funded by National Institutes of Health Grants to HNA (F31 DK121484), BJK (R01 DK115478), and LAL (F32 DK128969), and the Intramural 6 Research Program at the NIH National Center for Complementary and Integrative Health (YC).

## Author contributions

BJK and HNA designed all experiments. HNA, SC, VMH, and LAL completed all experiments. HNA, BJK, SC, and YC analyzed data. HNA wrote the manuscript with feedback and edits from BJK, YC, and BKT.

## References

1. Ocklenburg S, Güntürkün O (2017): The Lateralized Brain: The Neuroscience and Evolution of Hemispheric Asymmetries. The Lateralized Brain: The Neuroscience and Evolution of Hemispheric Asymmetries. Elsevier Inc. https://doi.org/10.1016/C2014-0-03755-0

2. Rogers LJ, Vallortigara G (2013): Divided Brains: The Biology and Behavior of Brain Asymmetries. Cambridge University Press.

3. Güntürkün O, Ströckens F, Ocklenburg S (2020): Brain lateralization: A comparative perspective. Physiol Rev 100: 1019–1063.

4. Bach DR, Herdener M, Grandjean D, Sander D, Seifritz E, Strik WK (2009): Altered lateralisation of emotional prosody processing in schizophrenia. Schizophr Res 110: 180–187.

5. Taylor SF, Liberzon I, Decker LR, Koeppe RA (2002): A functional anatomic study of emotion in schizophrenia. Schizophr Res. https://doi.org/10.1016/S0920-9964(01)00403-0

6. Phan KL, Orlichenko A, Boyd E, Angstadt M, Coccaro EF, Liberzon I, Arfanakis K (2009): Preliminary Evidence of White Matter Abnormality in the Uncinate Fasciculus in Generalized Social Anxiety Disorder. Biol Psychiatry 66: 691–694.

7. Hahn A, Stein P, Windischberger C, Weissenbacher A, Spindelegger C, Moser E, et al. (2011): Reduced resting-state functional connectivity between amygdala and orbitofrontal cortex in social anxiety disorder. Neuroimage. https://doi.org/10.1016/j.neuroimage.2011.02.064

8. Bruder GE, Stewart JW, Towey JP, Friedman D, Tenke CE, Voglmaier MM, et al. (1992): Abnormal cerebral laterality in bipolar depression: Convergence of behavioral and brain event-related potential findings. Biol Psychiatry 32: 33–47.

9. Farahbod H, Cook IA, Korb AS, Hunter AM, Leuchter AF (2010): Amygdala lateralization at rest and during viewing of neutral faces in major depressive disorder using low-resolution brain electromagnetic tomography. Clin EEG Neurosci 41: 19–23.

10. Frodl T, Meisenzahl E, Zetzsche T, Bottlender R, Born C, Groll C, et al. (2002): Enlargement of the amygdala in patients with a first episode of major depression. Biol Psychiatry 51: 708–14.

11. Rauch SL, Whalen PJ, Shin LM, Mcinerney SC, Macklin ML, Lasko NB, et al. (2000): Exaggerated Amygdala Response to Masked Facial Stimuli in Posttraumatic Stress Disorder: A Functional MRI Study. Biological Psychiatry 47: 769–776.

12. Symonds LL, Gordon NS, Bixby JC, Mande MM (2006): Right-lateralized pain processing in the human cortex: An fMRI study. J Neurophysiol 95: 3823–3830.

13. Allen HN, Bobnar HJ, Kolber BJ (2021, January 1): Left and right hemispheric lateralization of the amygdala in pain. Progress in Neurobiology, vol. 196. Elsevier Ltd. https://doi.org/10.1016/j.pneurobio.2020.101891

14. International Association for the Study of Pain (2003): How Prevalent is Chronic Pain? PAIN Clinical Updates XI.

15. Miller LR, Cano A (2009): Comorbid Chronic Pain and Depression: Who Is at Risk? Journal of Pain 10: 619–627.

16. Schmidt-Wilcke T (2015, February 1): Neuroimaging of chronic pain. Best Practice and Research: Clinical Rheumatology, vol. 29. Bailliere Tindall Ltd, pp 29–41.

17. Simons LE, Moulton EA, Linnman C, Carpino E, Becerra L, Borsook D (2014): The human amygdala and pain: Evidence from neuroimaging. Hum Brain Mapp 35: 527–538.

18. Neugebauer V, Li W, Bird GC, Han JS (2004): The amygdala and persistent pain. Neuroscientist 10: 221–234.

19. Baas D, Aleman A, Kahn RS (2004): Lateralization of amygdala activation: A systematic review of functional neuroimaging studies. Brain Res Rev 45: 96–103.

20. Carrasquillo Y, Gereau IV RW (2008): Hemispheric lateralization of a molecular signal for pain modulation in the amygdala. Mol Pain 4. https://doi.org/10.1186/1744-8069-4-24

21. Ji G, Neugebauer V (2009): Hemispheric Lateralization of Pain Processing by Amygdala Neurons. J Neurophysiol 102: 2253–2264.

22. Wartolowska K, Hough MG, Jenkinson M, Andersson J, Wordsworth BP, Tracey I (2012): Structural changes of the brain in rheumatoid arthritis. Arthritis Rheum 64: 371–379.

23. Kulkarni B, Bentley DE, Elliott R, Youell P, Watson A, Derbyshire SWG, et al. (2005): Attention to pain localization and unpleasantness discriminates the functions of the medial and lateral pain systems. European Journal of Neuroscience 21: 3133–3142.

24. Sadler KE, McQuaid NA, Cox AC, Behun MN, Trouten AM, Kolber BJ (2017): Divergent functions of the left and right central amygdala in visceral nociception. Pain 158: 747–759.

25. Miyazawa Y, Takahashi Y, Watabe AM, Kato F (2018): Predominant synaptic potentiation and activation in the right central amygdala are independent of bilateral parabrachial activation in the hemilateral trigeminal inflammatory pain model of rats. Mol Pain 14. https://doi.org/10.1177/1744806918807102

26. Kolber BJ, Montana MC, Carrasquillo Y, Xu J, Heinemann SF, Muglia LJ, Gereau RW (2010): Activation of Metabotropic Glutamate Receptor 5 in the Amygdala Modulates Pain-Like Behavior. Journal of Neuroscience 30: 8203–8213.

27. Carrasquillo Y, Gereau RW (2007): Activation of the Extracellular Signal-Regulated Kinase in the Amygdala Modulates Pain Perception. Journal of Neuroscience 27: 1543–1551.

28. Ikeda R, Takahashi Y, Inoue K, Kato F (2007): NMDA receptor-independent synaptic plasticity in the central amygdala in the rat model of neuropathic pain. Pain. https://doi.org/10.1016/j.pain.2006.09.003

29. Cooper AH, Brightwell JJ, Hedden NS, Taylor BK (2018): The left central nucleus of the amygdala contributes to mechanical allodynia and hyperalgesia following right-sided peripheral nerve injury. Neurosci Lett. https://doi.org/10.1016/j.neulet.2018.08.013

30. Wall PD, Melzack R, Bonica JJ (1999): Textbook of Pain. Churchill Livingston Edinburgh.

31. Shoskes DA, Nickel JC, Rackley RR, Pontari MA (2009): Clinical phenotyping in chronic prostatitis/chronic pelvic pain syndrome and interstitial cystitis: A management strategy for urologic chronic pelvic pain syndromes. Prostate Cancer Prostatic Dis 12: 177–183.

32. Adamian L, Urits I, Orhurhu V, Hoyt D, Driessen R, Freeman JA, et al. (2020, June 1): A Comprehensive Review of the Diagnosis, Treatment, and Management of Urologic Chronic Pelvic Pain Syndrome. Current Pain and Headache Reports, vol. 24. Springer. https://doi.org/10.1007/s11916-020-00857-9

33. Zhang J, Liang CZ, Shang X, Li H (2020, January 1): Chronic Prostatitis/Chronic Pelvic Pain Syndrome: A Disease or Symptom? Current Perspectives on Diagnosis, Treatment, and Prognosis. American Journal of Men’s Health, vol. 14. SAGE Publications Inc. https://doi.org/10.1177/1557988320903200

34. Boezaart AP, Smith CR, Chembrovich S, Zasimovich Y, Server A, Morgan G, et al. (2021): Visceral versus somatic pain: an educational review of anatomy and clinical implications. Reg Anesth Pain Med 46: 629–636.

35. As-Sanie S, Harris RE, Napadow V, Kim J, Neshewat G, Kairys A, et al. (2012): Changes in regional gray matter volume in women with chronic pelvic pain: A voxel-based morphometry study. Pain 153: 1006–1014.

36. Bagarinao E, Johnson KA, Martucci KT, Ichesco E, Farmer MA, Labus J, et al. (2014): Preliminary structural MRI based brain classification of chronic pelvic pain: A MAPP network study. Pain 155: 2502–2509.

37. Kleinhans NM, Yang CC, Strachan ED, Buchwald DS, Maravilla KR (2016): Alterations in Connectivity on Functional Magnetic Resonance Imaging with Provocation of Lower Urinary Tract Symptoms: A MAPP Research Network Feasibility Study of Urological Chronic Pelvic Pain Syndromes. Journal of Urology 195: 639–645.

38. Nation KM, DeFelice M, Hernandez PI, Dodick DW, Neugebauer V, Navratilova E, Porreca F (2018): Lateralized Kappa Opioid Receptor Signaling from the Amygdala Central Nucleus Promotes Stress-Induced Functional Pain. Pain 159: 1.

39. Andreoli M, Marketkar T, Dimitrov E (2017): Contribution of amygdala CRF neurons to chronic pain. Exp Neurol. https://doi.org/10.1016/j.expneurol.2017.08.010

40. Wilson TD, Valdivia S, Khan A, Ahn H-S, Adke AP, Gonzalez SM, et al. (2019): Dual and Opposing Functions of the Central Amygdala in the Modulation of Pain. Cell Rep 29: 332-346.e5.

41. Bourgeais L, Gauriau C, Bernard JF (2001): Projections from the nociceptive area of the central nucleus of the amygdala to the forebrain: A PHA-L study in the rat. European Journal of Neuroscience 14: 229–255.

42. Chiang MC, Bowen A, Schier LA, Tupone D, Uddin O, Heinricher MM (2019): Parabrachial complex: A hub for pain and aversion. Journal of Neuroscience 39: 8225–8230.

43. Sun L, Liu R, Guo F, Wen M qing, Ma X lin, Li K yuan, et al. (2020): Parabrachial nucleus circuit governs neuropathic pain-like behavior. Nat Commun 11. https://doi.org/10.1038/s41467-020-19767-w

44. Schwaber JS, Sternini C, Brecha NC, Rogers WT, Card JP (1988): Neurons Containing Calcitonin Gene-Related Peptide in the Parabrachial Nucleus Project to the Central Nucleus of the Amygdala. Journal of Comparative Neurology 270: 416–426.

45. Palmiter RD (2018, May 1): The Parabrachial Nucleus: CGRP Neurons Function as a General Alarm. Trends in Neurosciences, vol. 41. Elsevier Ltd, pp 280–293.

46. Boudes M, Uvin P, Kerselaers S, Vennekens R, Voets T, De Ridder D (2011): Functional characterization of a chronic cyclophosphamide-induced overactive bladder model in mice. Neurourol Urodyn 30: 1659–1665.

47. Stillwell TJ, Benson RC (1988): Cyclophosphamide-induced hemorrhagic cystitis: A review of 100 patients. Cancer 61: 451–457.

48. Sadler KE, Stratton JM, Kolber BJ (2014): Urinary bladder distention evoked visceromotor responses as a model for bladder pain in mice. Journal of Visualized Experiments. https://doi.org/10.3791/51413

49. Tye KM, Prakash R, Kim SY, Fenno LE, Grosenick L, Zarabi H, et al. (2011): Amygdala circuitry mediating reversible and bidirectional control of anxiety. Nature 471: 358–362.

50. Chaplan SR, Bach FW, Pogrel JW, Chung JM, Yaksh TL (1994): Quantitative assessment of tactile allodynia in the rat paw. Journal of Neuroscience Methods, vol. 53.

51. Block CH, Hoffman G, Kapp BS (1989): Peptide-containing pathways from the parabrachial complex to the central nucleus of the amygdala. Peptides, vol. 10. https://doi.org/10.1016/0196-9781(89)90060-0

52. Sugimura YK, Takahashi Y, Watabe AM, Kato F (2016): Synaptic and network consequences of monosynaptic nociceptive inputs of parabrachial nucleus origin in the central amygdala. J Neurophysiol 115: 2721–2739.

53. Felix-Ortiz AC, Beyeler A, Seo C, Leppla CA, Wildes CP, Tye KM (2013): BLA to vHPC inputs modulate anxiety-related behaviors. Neuron 79: 658–664.

54. Kim J, Zhang X, Muralidhar S, LeBlanc SA, Tonegawa S (2017): Basolateral to Central Amygdala Neural Circuits for Appetitive Behaviors. Neuron 93: 1464-1479.e5.

55. Shinohara K, Watabe AM, Nagase M, Okutsu Y, Takahashi Y, Kurihara H, Kato F (2017): Essential role of endogenous calcitonin gene-related peptide in pain-associated plasticity in the central amygdala. European Journal of Neuroscience. https://doi.org/10.1111/ejn.13662

56. Han JS, Li W, Neugebauer V (2005): Critical Role of Calcitonin Gene-Related Peptide 1 Receptors in the Amygdala in Synaptic Plasticity and Pain Behavior. The Journal of Neuroscience 25: 10717–10728.

57. Han JS, Adwanikar H, Li Z, Ji G, Neugebauer V (2010): Facilitation of synaptic transmission and pain responses by CGRP in the amygdala of normal rats. Mol Pain 6. https://doi.org/10.1186/1744-8069-6-10

58. Okutsu Y, Takahashi Y, Nagase M, Shinohara K, Ikeda R, Kato F (2017): Potentiation of NMDA receptormediated synaptic transmission at the parabrachial-central amygdala synapses by CGRP in mice. Mol Pain 13: 1–11.

59. Sugimoto M, Takahashi Y, Sugimura YK, Tokunaga R, Yajima M, Kato F (2021): Active role of the central amygdala in widespread mechanical sensitization in rats with facial inflammatory pain. Pain. https://doi.org/10.1097/j.pain.0000000000002224

60. Xu W, Lundeberg T, Wang YT, Li Y, Yu LC (2003): Antinociceptive effect of calcitonin gene-related peptide in the central nucleus of amygdala: Activating opioid receptors through amygdala-periaqueductal gray pathway. Neuroscience 118: 1015–1022.

61. Butler RK, Ehling S, Barbar M, Thomas J, Hughes MA, Smith CE, et al. (2017): Distinct neuronal populations in the basolateral and central amygdala are activated with acute pain, conditioned fear, and fearconditioned analgesia. Neurosci Lett 661: 11–17.

62. Ji G, Neugebauer V (2007): Differential effects of CRF1 and CRF2 receptor antagonists on pain-related sensitization of neurons in the central nucleus of the amygdala. J Neurophysiol 97: 3893–3904.

63. Neugebauer V, Mazzitelli M, Cragg B, Ji G, Navratilova E, Porreca F (2020): Amygdala, neuropeptides, and chronic pain-related affective behaviors. Neuropharmacology 108052.

64. Gonçalves L, Dickenson AH (2012): Asymmetric time-dependent activation of right central amygdala neurones in rats with peripheral neuropathy and pregabalin modulation. European Journal of Neuroscience 36: 3204–3213.

65. As-Sanie S, Harris RE, Napadow V, Kim J, Neshewat G, Kairys A, et al. (2012): Changes in regional gray matter volume in women with chronic pelvic pain: A voxel-based morphometry study. Pain 153: 1006–1014.

66. Chiang MC, Nguyen EK, Canto-Bustos M, Papale AE, Oswald AMM, Ross SE (2020): Divergent Neural Pathways Emanating from the Lateral Parabrachial Nucleus Mediate Distinct Components of the Pain Response. Neuron 106: 927-939.e5.

67. Han S, Soleiman M, Soden M, Zweifel L, Palmiter RD (2015): Elucidating an Affective Pain Circuit that Creates a Threat Memory. Cell 162: 363–374.

68. Corder G, Ahanonu B, Grewe BF, Wang D, Schnitzer MJ, Scherrer G (2019): An amygdalar neural ensemble that encodes the unpleasantness of pain. Science, vol. 363. https://doi.org/10.1126/science.aap8586

