## Supplemental Information for "A parabrachial-to-amygdala circuit that determines hemispheric lateralization of somatosensory processing"

**Supplemental Methods**

**Animals**

All experiments used *Calca^tm1.1(cre/EGFP)Rpa^* (*Calca^Cre^*) Cre-recombinase knockin mice (Jax033168; originally obtained from Dr. Richard Palmiter, University of Washington), *Calcrl*^Cre^ (obtained from Dr. Richard Palmiter, University of Washington) crossed with a *Rosa26-flox-stop-tdTomato* reporter line Ai9 (Jax Ai9- 007909) to generate *Calcrl*^Cre^::Ai9, or wild-type C57BL/6J (Jax000664) female littermates aged 9-13 weeks of age at the time of experimentation. Wild type breeders were acquired from Jackson Laboratory and replaced every 4 generations. *Calca^Cre^* (1) and Calcrl^Cre^ (2) mice were bred as heterozygous pairs and backcrossed to C57BL/6J mice from Jackson Labs. Genotypes were confirmed using PCR, validating the expression of the wild-type allele and transgene in Cre heterozygous animals and the lack of the wild-type allele in Cre homozygous (“CGRP-knockout”) animals. Immunohistochemistry for CGRP was also used to confirm *Calca* knockout in *Calca^Cre/Cre^* homozygous animals. Transgenic animals were bred in house at Duquesne University (Pittsburgh, PA) and housed 2-5 animals per cage on a 12-h light/dark schedule (7 am-7 pm) with *ad libitum* access to food and water. For electrophysiology studies at the National Institutes of Health and behavioral studies at the University of Texas at Dallas, *Calcrl*^Cre^::Ai9 and C57BL/6J mice were bred in-house. Behavior and physiology assays were conducted during the light cycle. All experiments were in accordance with the National Institutes of Health guidelines and were approved by the Animal Care and Use Committee at Duquesne University (protocols 1905-06 and 2006-03), University of Pittsburgh (protocol 21063864), University of Texas at Dallas (protocol 20-04) or the National Institutes of Health (protocol 1397).

**Cyclophosphamide-induced cystitis**

Cyclophosphamide (CYP) is a chemotherapeutic drug which is used experimentally broken into various metabolites including acrolein which causes inflammation of bladder tissue and a bladder pain-like sensitivity phenotype in rodents (3,4). Animals receive three 100 μL intraperitoneal injections of 100 mg/kg CYP (Sigma) every other day for five days prior to experimentation. CYP solutions were made fresh on the day of the injection. Control mice received 100 μL intraperitoneal injections of saline. All experiments were completed one day after the completion of the CYP protocol (day 6).

**Stereotaxic surgeries**

*Viral injection*

Adeno-associated viruses containing Cre-dependent optogenetic constructs (rAAV5/Ef1a-DIO-hCHR2-mCherry, rAAV5/EF1a-DIO-NpHR3.0-mCherry, or rAAV5/EF1a-DIO-mCherry) were used to manipulate activity of *Calca*-expressing fibers from the parabrachial nucleus (PBN) in the central amygdala (CeA). *Calca^Cre+/-^* mice 6-8 weeks old were anesthetized under 2% isoflurane and placed in a stereotaxic frame; body temperature was maintained with a heating pad throughout the surgery. A craniotomy was performed over either the left, right, or bilateral PBN and 1 μL of virus was injected into the PBN (AP -5.15 mm; ML +/-1.45 mm, DV -3.45 mm) via a 2.5 μL Hamilton syringe connected to a glass pipette at a rate of 0.20μL/min using the Quintessential Stereotax Injector (Stoelting). The pipette was left in place for an additional five minutes to allow for diffusion. The experimenter was blinded to virus being injected.

*Cannula/wireless LED implantation*

For cannula implantation following viral injection, mice received a craniotomy over the left, right, or bilateral CeA ipsilateral to viral injection as previously described (5). Two bone screws (Stoelting) were implanted on either side of the CeA craniotomy, anterior to bregma and anterior to lambda. A stainless-steel cannula (8.01 mm long, 0.2 mm diameter) was lowered into the CeA (AP -1.45 mm, ML +/- 3.00 mm, DV -4.20 mm) and fixed in place using dental cement. Animals receiving a 4-mm wireless LED Neurolux device (Neurolux, St. Louis, MO) following viral injection had the device lowered into the CeA ipsilateral to PBN viral injection and fixed in place using superglue. The scalp was sutured over the device. Animals recovered from anesthesia on a heating pad before being returned to their homecage. Animals received topical lidocaine on scalp incisions and had free access to children’s Motrin (40mg/kg/day in drinking water) for 72 hours post-surgery. Behavioral experiments were conducted 3 weeks after surgery to allow for optimal viral expression and full surgical recovery.

For animals receiving only cannulae (pharmacology experiments, ELISA experiment), a craniotomy was performed over the left, right, or bilateral CeA. In control placement experiments, cannulae were implanted over the striatum (AP -1.45 mm, M/L +/- 3.00 mm, D/V -3.50 mm). Bone screws (Stoelting) were implanted contralateral to the side of cannulation anterior to bregma and anterior to lambda. A stainless-steel cannula was lowered into the left, right, or bilateral CeA (or striatum) and fixed in place with dental cement (Stoelting). Animals were allowed to recover from anesthesia on a heating pad before being returned to their homecage. Animals received topical lidocaine on scalp incisions and had free access to children’s Motrin (40 mg/kg/day in drinking water) for 72 hours post-surgery. Behavioral experiments were conducted 1-2 weeks following cannulation.

**Urinary bladder distention**

Urinary bladder distention (UBD) was conducted in 9- to 13-week-old female mice and visceromotor responses (VMR) were recorded by measuring electromyography (EMG) of the external abdominal oblique muscle during noxious distention as a quantitative measure of bladder pain-like responses. UBD-VMR was performed as previously described (6) one day following the final injection of CYP (day 6). Mice were anesthetized using isoflurane in an induction chamber before being transferred to a nose cone with 2% isoflurane vaporized in 100% oxygen. A lubricated 24-guage, 14 mm catheter was inserted into the bladder via the urethra. We have previously shown that the side of body recorded does not impact brain lateralized responses (5). Thus, the left abdominal wall was exposed, and two silver wires were implanted in the left external abdominal oblique muscle. A third wire was passed through the skin of the chest to serve as a grounding wire. Body temperature was maintained at 37^o^C throughout the experiment using a battery-operated heating pad as well as an overhead radiant heat lamp.

Following surgery, isoflurane was lowered to 1.5% and then further lowered in 0.125% steps every 10 minutes until the animal responded to a noxious toe pinch but did not ambulate of vocalize (approximately 0.875-1.0% isoflurane). Once a stable level of isoflurane was reached, animals’ bladders were distended 5-10 times with 60 mmHg of compressed air administered using a custom timed-pressure regulator (Washington University School of Medicine, St. Louis, MO) to establish stable VMRs to distention. Each distention lasted 20 seconds with a 1-minute intertrial interval between distentions. EMG signals were relayed via an amplifier through a Cambridge Electronic Design (CED) 1401 module to a computer with Spike2 software. Data was exported to IgorPro where background EMG was subtracted and stimulus evoked EMGs were rectified and integrated over the 20 second pressure period using a custom script. A second similar custom script was used to analyze the pre-distention period, and all VMRs were normalized to the smallest pre-distention VMR.

*Optogenetic manipulation during UBD*

Following the establishment of stable VMRs to bladder distention, baseline VMRs were collected (10 distentions alternating at 30 mmHg and 60 mmHg pressure). Immediately after baseline, light (532 nm for inhibition or 473 nm for excitation) was delivered using a low-power laser diode (Shanghai Laser and Optics Century Co, Ltd, Shanghai, China) or LED (Thor-Labs, Newton, NJ) during the “light-on” timepoint and VMRs were collected (same parameters as baseline). Immediately after the completion of the “light-on” timepoint, the light source was turned off and the post light timepoint was collected using the same pressure sequence parameters.

For inhibition experiments, 532 nm constant light stimulation (8-15 mW power) was administered. For excitation experiments, 473 nm light was pulsed (20 Hz, 5-ms pulse width, 3-20 mW power) to mimic the firing of CeA neurons following injury (7). Experimenter was blinded to virus (mCherry v ChR2/NpHR) until optogenetic manipulation was completed.

**Pharmacological activation/inhibition during UBD**

CGRP and CGRP(8-37) were purchased from Genscript (RP11095, RP11090) and reconstituted and diluted in aCSF. Following the collection of baseline VMRs, wild type animals received injection of aCSF, 100 nM CGRP (volume 1 μL), 100 nM CGRP(8-37), or a cocktail of 100 nM CGRP+100 nM CGRP(8-37) at a rate of 0.2 μL/min via a 32-gauge injection cannula that extended 0.1 mm beyond the tip of the cannula. The injection cannula was coupled to a Hamilton syringe via flexible plastic tubing, and the injector was left in place for an additional 5 minutes to allow for diffusion. VMRs were collected in response to UBD using the same parameters as baseline every fifteen minutes following the completion of injection for 90 minutes. Immediately following the completion of the experiment, animals were euthanized, and brains were extracted for cannula placement verification. The experimenter was blinded to drug treatment until after verification of post hoc viral expression and/or cannula placement targeting verification. For striatum placement control experiment, experimenter was blinded to experimental hypothesis, cannula placement, and treatment but not side of brain.

**Combined optogenetic and pharmacological manipulation during UBD**

Baseline VMRs were collected as in other UBD experiments in *Calca^Cre^* mice. Immediately following baseline, 1 μL of aCSF, 100 nM CGRP(8-37), or a cocktail of 22 mM AP5 and 38 mM NBQX (8) was injected into either the left or right CeA as described above. Thirty minutes following injection, a fiberoptic coupled to a 473-nm laser was inserted into the cannula and pulsed at 20-Hz (15-20-mW) while VMRs were collected. Post laser and post drug VMRs were collected 90 min following injection to allow time for drug washout. Experimenter was blinded to drug injection until viral injection site and cannula placement were verified.

For *Calca^Cre/+^* heterozygote versus knockout (*Calca^Cre/Cre^*) experiments, baseline VMRs were collected as described. Following baseline, animals had either a fiberoptic coupled to a 473-nm laser for optogenetic excitation or an injector for pharmacology inserted into the cannula. Optogenetic and pharmacology experiments were performed in the same animals in a randomized order and no effect of order was found (**Supplementary Fig. 7**). Optogenetic activation of CGRP terminals in the left or right CeA was performed identical to optogenetic manipulation during UBD described above. Pharmacological activation of CGRP receptor cells in the CeA was achieved via infusion of CGRP as described above. Experimenter was blinded to genotype until viral injection site and cannula placement were verified.

**von Frey abdominal sensitivity**

*In vivo* behavioral testing was conducted one to two days following final CYP injection (day 6-7). This correlates to day 20 in experiments where animals received optogenetic stimulation of CGRP-containing PBN fibers. Animals were habituated in ventilated Plexiglas enclosures 15 cm x 15 cm on wire mesh for 1.5 hours before testing. Sixty decibel white noise was used to mask background sounds. The experimenter was present in the room for 30 min prior to the beginning of testing. Calibrated von Frey filaments (Touch Test) were used to assess abdominal sensitivity on the right and left abdomen approximately 0.5 cm from the urethra via the up-down method to calculate 50% withdrawal thresholds (9). No difference was found in abdominal sensitivity between right and left side, so withdrawal thresholds were averaged.

*Optogenetic manipulation during abdominal sensitivity testing*

Animals were habituated on metal mesh in Plexiglas boxes tuned for Neurolux optogenetic stimulation. Following collection of baseline 50% withdrawal thresholds, a 473-nm wireless LED Neurolux device was remotely activated (20-Hz stimulation, 5-ms pulse width, 10mW power) and 50% withdrawal threshold were collected again. Experimenter was blinded to virus (mCherry v ChR2) until after analysis of viral injection site and LED placement.

**Real time place preference**

One day prior to abdominal von Frey testing (day 6 post initial CYP injection, day 19 post-surgery), animals were habituated to the behavior room for 20 min in their home cage. Animals were habituated on day 6 to avoid a negative association of CYP injection with habituation. Sixty decibel white noise was used to mask background noise. Animals were placed in a three-chamber Plexiglas place preference apparatus (30 cm^2^ x 20 cm) with distinct visual patterns and allowed to freely explore for 20 min. The animals’ activity was video recorded using AnyMaze (Stoelting Co.) behavioral tracking software. The next day (day 20 post-surgery), following von Frey abdominal sensitivity testing, animals were returned to their home cage for 20 min. Animals were then placed back in the place preference apparatus where one chamber was tuned for wireless Neurolux LED stimulation. Upon entering the tuned chamber, the Neurolux device automatically started stimulation (473-nm, 20-Hz, 10mW power), which ended as soon as the animal exited the Neurolux tuned chamber. Animals’ activity was video recorded for 20 min using AnyMaze. Any animal displaying more than 700 seconds in a single chamber on habituation day was excluded from the experiment. Four animals were excluded. Experimenter was blinded to virus (mCherry v ChR2) until after analysis of viral injection site and LED placement.

**Immunohistochemistry**

*Viral Targeting*

All viral constructs used in these experiments contained an mCherry sequence to allow for viral targeting in *Calca-*expressing (Cre-recombinase positive) cells in the PBN and terminals in the CeA. Following optogenetic experiments, animals were perfused using 20 mL 1x phosphate buffered saline (PBS) and 20 mL ice cold 4% paraformaldehyde (PFA). Brains were removed and stored in 4% PFA overnight at 4°C before being transferred to 30% sucrose. After 3-5 days in sucrose, brains were frozen and kept at -80°C until sectioning. Thirty µm coronal sections were collected using a cryostat and stored in PBS at 4^o^C. Immunohistochemistry was performed on floating PBN and CeA sections for mCherry. Sections were washed 3 times in 1x PBS, blocked for 60 minutes using 1% bovine serum albumin and 0.2% milk in 1x PBS with 0.1% Triton-X, and incubated overnight at 4^o^C in 1:1000 anti-mCherry (rabbit, Abcam ab167453) in blocking solution. Sections were then washed 3 times in PBS with 0.1% Triton X before being incubated in 1:1000 anti-555 (anti-rabbit Alexa Fluor 555, Invitrogen A-21249) in blocking solution at room temperature for 1 hour. Sections were then washed 3 times in PBS before being mounted onto Superfrost Plus (Fisher Scientific) microscope slides, coverslipped with Vectashield (Vector Laboratories) anti-fade mounting medium with DAPI, and imaged using a Nikon Eclipse Ti2 microscope.

*CGRP quantification*

C57BL/6J female mice were treated with either CYP or saline and perfused one day following the final injection (day 6). Thirty µm coronal sections were collected and stored in PBS at 4 ^o^C until staining. Six representative sections from across the rostral-caudal axis of the CeA were picked for staining. Tissue was washed three times in 1x PBS before being blocked in 10% normal goat serum in PBS with 0.1% Triton-X for 60 minutes. Tissue was incubated with anti-CGRP (1:2000 rabbit anti CGRP, Calbiochem PC205L) in 5% normal goat serum in PBS with 0.1% Triton-X overnight at 4 ^o^C. Tissue was then washed three times in 1x PBS with 0.1% Triton-X and incubated in fluorescent secondary antibody (goat anti-rabbit AlexaFluor 488, Life Technologies, A-11034) diluted in 5% normal goat serum in 1x PBS with 0.1% Triton-X for 60 minutes at room temperature. After three more washes in 1x PBS, tissue sections were mounted on SuperFrost Plus microscope slides and coverslipped with Vectashield anti-fade mounting medium with DAPI. Images were captured using a Nikon confocal microscope and fluorescence intensity for each channel was quantified using NIS-Elements Advanced Research software. All microscope images were acquired using settings from a negative control and settings were kept consistent. Fluorescence intensity of the 488 channel was normalized to fluorescence intensity of the DAPI channel for each image. The CeC was defined as the area 200 µm inward from BLA/CeA border (24).

***Ex-vivo* Electrophysiology**

*Acute slice preparation of left and right CeA*

Female *Calcrl*^Cre^::Ai9 or C57BL/6J wild-type mice (9 to 13 weeks old) were deeply anesthetized with 1.25% Avertin (0.4 mg/g) injected intraperitoneally and transcardially perfused with ice-cold cutting solution (110 mM choline chloride, 25 mM NaHCO_3_, 1.25 mM NaH_2_PO_4_, 2.5 mM KCl, 0.5 mM CaCl_2_, 7.2 mM MgCl_2_, 25 mM D-glucose,12.7 mM L-ascorbic acid, 3.1 mM pyruvic acid, oxygenated with 95%/5% O_2_/CO_2_). Brains were extracted, placed in ice-cold cutting solution, and cut in coronal slices (250 μm) with a Leica VT1200 S vibrating blade microtome (Leica Microsystems Inc). CeA slices were cut in half to separate left and right hemispheres and were incubated at 33°C for 30 minutes in artificial cerebrospinal fluid (aCSF) (125 mM NaCl, 2.5 mM KCl, 1.25 mM NaH_2_PO_4_, 25 mM NaHCO_3_, 2 mM CaCl_2_, 1 mM MgCl2, 25 mM D-glucose). Slices were recovered at room temperature for at least 20 min before recording. Chambers were continuously oxygenated with 95%/5% O_2_/CO_2_ during incubation and recovery.

*Electrophysiological recordings*

The recording chamber was perfused continuously with aCSF oxygenated with 95%/5% O_2_/CO_2_ (1ml/min), and all recordings were performed at 33 ± 1^°^C using a recording chamber heater and in-line solution heater (Warner Instruments). Recording pipettes (2-6MΩ resistance) were filled with internal solution (120mM potassium methyl sulfate, 20 mM KCl, 10 mM HEPES, 0.2 mM EGTA, 8 mM NaCl_2_, 4 mM Mg-ATP, 0.3 mM Tris-GTP, and 14 mM phosphocreatine with pH 7.3 using 5 M KOH and an osmolarity of ~300 mosmol^-1^). Whole-cell current clamp recordings were collected from neurons in the capsular (CeC) subdivision of either the right or left CeA, as identified using an upright microscope (Nikon Eclipse FN1). Recordings were acquired using the Multiclamp 700B patch-clamp amplifier interfaced with a Digidata 1500 acquisition system and pCLAMP 10.7 software (Molecular Devices) on a Dell computer. Before forming a membrane-pipette seal, pipette tip potentials were zeroes. Series resistances (not exceeding 20 MΩ) were monitored throughout the recordings. Whole-cell capacitance was measured in voltage-clamp configuration, with the cell held at -70 mV, then subjected to a +/- 10 mV current change of 25 ms duration.

Recordings were restricted to late-firing fluorescently labeled neurons expressing the CGRP receptor (CGRPR+) in slices from *Calcrl*^Cre^::Ai9 mice or unlabeled neurons in slices from C57BL/6J wild-type mice within the capsular subdivision of the CeA. Late-firing neurons were defined as cells with a latency to spike higher than 100 ms at a 280 pA current amplitude as previously described (10). 500ms depolarizing current of various amplitudes (between 20 pA and 360 pA) was injected from resting membrane potential to elicit repetitive action potentials firing under an initial aCSF bath condition and after bath application of 500 nM CGRP (11,12) for at least 4 min to allow for a full bath exchange. Only 1 cell per slice was used following CGRP bath application. Recording from control cells were obtained before and 4 min after bath exchange to aCSF. Recordings were acquired at 100 kHz and filtered at 10 kHz. At the end of each electrophysiological recording, pipette location was imaged at low magnification and the anatomical localization of each recording was determined in reference to a mouse brain atlas (13).

**RNAscope *in situ* hybridization**

RNAscope fluorescent multiplex assay v2 was used with probes for *Calcrl*, *Prkcd,* and *Sst* to determine co-localization in the CeA. Brains from perfused C57Bl/6J mice treated with either CYP or saline were postfixed in 4% PFA for 3 hours before being transferred to 30% sucrose until tissue sank. Brains were then frozen and 20 μm coronal sections were collected using a cryostat and stored in anti-freeze at -20^o^C. Five to six representative sections from across the rostral-caudal axis of the CeA in each animal were mounted on Superfrost Plus slides and allowed to air dry overnight at room temperature. Slides were treated twice with xylene (5 minutes each) followed by two treatments with 100% ethanol (2 min each). Tissue was treated with protease III for 20 min at 40^o^C in HybEZ oven. RNAscope was performed according to manufacturer’s instructions (ACDBio Inc.). Briefly, probes for *Calcrl,* *Prkcd*, and *Sst* were hybridized for 2 hours in the HybEZ oven at 40^o^C. Signal was amplified using a series of AMPs according to manufacturer’s instructions, followed by development of a TSA-based fluorescent label. Slides were cover slipped with Vectashield anti-fade mounting medium with DAPI. Images were captured within 48 hours using a Nikon Eclipse Ti2 microscope and images were analyzed using NIS-Elements Advanced Research software. Positive cells were identified as a DAPI-labeled nucleus surrounded by at least three puncta. Cell counts were determined blinded to treatment (CYP vs saline). Cell number and percent co-localization were averaged across all sections from the same brain.

**CeA tissue collection and cAMP ELISA**

Wild-type C57Bl6/J mice with bilateral cannulae were anesthetized using isoflurane in an induction chamber before being transferred to a nose cone with 1% isoflurane vaporized in 100% oxygen. CGRP purchased from Genscript (RP11095, RP11090) was reconstituted and diluted to 100nM in aCSF. Animals received bilateral injections of aCSF or 100 nM CGRP (1 μL) at a rate of 0.2 μL/min via a 32-gauge injection cannula that extended 0.1 mm beyond the tip of the cannula. The injection cannula was coupled to a Hamilton syringe via flexible plastic tubing, and the injector was left in place for an additional 5 minutes to allow for diffusion. Following infusion, mice were returned to their homecage for 40 minutes. Mice were decapitated and brains were sectioned and flash frozen to extract CeA micro punches. Left and right CeA tissue from each animal was homogenized in 1X homogenate buffer (20 mM Tris pH 7.5, 1 mM EDTA, 1 mM Na_4_P_2_O_7_, 1X Halt Thermo Scientific Protease Inhibitor Single-Use Cocktail, PI78425) and centrifuged at 12,000 RPM for 20 minutes at 4^o^C. Supernatant was collected and stored at -80^o^C until ELISA was performed. ELISA was performed according to kit instructions (Cell Biolabs, STA-500) using tissue homogenate from the left and right CeA.

**Statistics and data analysis**

All data analyses, including RNAscope, immunohistochemistry, behavior, and physiology, were conducted blind to treatment/virus/genotype. Data were analyzed using GraphPad Prism v9.1.2, NIS-Elements Advanced Research software v5.02, Spike2 v7.08, IgorPro v6.22A, AnyMaze (Version 9.0), and Microsoft Office v16.50. UBD data was analyzed via unpaired *t*-tests for two group comparisons, repeated-measures two-way analysis of variance (ANOVA) followed by Bonferroni or Dunnett’s post hoc tests for multiple comparisons. Behavioral data was analyzed using paired *t*-tests, one-way ANOVA, or repeated measures two-way ANOVAs followed by Bonferroni or Dunnett’s post hoc tests for multiple comparisons. *Ex vivo* physiology analysis was performed using Clampfit 10.7 software (Molecular Devices). Input resistance (R_in_) at resting membrane potential was calculated from the change in membrane potential resulting from a ± 20-pA current injection lasting 500 ms. Rheobase was defined as the minimal current required to evoke action potential activity from a 500-ms depolarizing current injection. Whole-cell membrane capacitance was calculated by integrating the capacitive transients elicited by a 25-ms voltage step (±10  mV) from −70  mV. Electrophysiology data were analyzed via two-way repeated measures ANOVA. RNAscope data was analyzed using two-way ANOVAs followed by Tukey post-hoc test for multiple comparisons. Statistical significance was determined at the level of P<0.05. Asterisks denoting P values include: *P<0.05, **P<0.01, ***P<0.001, and ****P<0.0001. All data are presented as the mean +/- standard error of the mean (SEM). Statistical information for all figures is provided in **Supplementary Table 1**.

**Supplementary Table 1**

| **Figure** | **Comparison** | **Analysis** | **P value, F, or t value** | **N** |
| --- | --- | --- | --- | --- |
| 1D | Normalized VMRs to 60 mmHg distention in naïve vs CYP- treated animals | Unpaired t-test | P=0.0076  t=2.776 | 28 mice per group |
| 1E | Percent change in VMR to 60 mmHg in CYP mice with mCherry or ChR2 in the left CeA before, during, and after laser stimulation | Two-way RM ANOVA with Dunnett’s posttest | Time, p=0.0025,  F (2, 36) = 7.132  Virus, p=0.0463,  F (1, 18) = 4.582  Time x virus, p=0.0018,  F (2, 36) = 7.543  mCherry:  BL v laser, p=0.9922  BL v PL, p=0.9205  ChR2:  BL v laser, p<0.0001  BL v PL, p>0.9999 | 10-11 mice per group |
| 1F | Area under the curve for 60 mmHg pressure in CYP mice with mCherry or ChR2 in the left CeA | Unpaired t-test | P=0.0076  t=3.007 | 10-11 mice per group |
| 1H | Percent change in VMR to 60 mmHg in CYP mice with mCherry or ChR2 in the right CeA before, during, and after laser stimulation | Two-way RM ANOVA with Dunnett’s posttest | Time, p=0.0059,  F (1.909, 30.55) = 6.242  Virus, p=0.2222  F (1, 16) = 1.613  Time x virus, p=0.0043,  F (2, 32) = 6.486  mCherry:  BL v laser, p=0.6944  BL v PL, p=0.8047  ChR2:  BL v laser, p=0.0071  BL v PL, p=0.4105 | 8-10 mice per group |
| 1I | Area under the curve for 60 mmHg pressure in CYP mice with mCherry or ChR2 in the right CeA | Unpaired t-test | P=0.0505  t=2.115 | 8-10 mice per group |
| 1K | Percent change in VMR to 60 mmHg in CYP mice with mCherry or NpHR in the left CeA before, during, and after laser stimulation | Two-way RM ANOVA with Dunnett’s posttest | Time, p=0.0171, F (2,2 2) = 4.923  Virus, p=0.0069, F (1, 11) = 10.98  Time x virus, p=0.0023, F (2, 22) = 8.078  mCherry:  BL v laser, p=0.7686  BL v PL, p=0.9967  NpHR:  BL v laser, p=0.0002  BL v PL, p=0.7854 | 6-7 mice per group |
| 1L | Area under the curve for 60 mmHg pressure in CYP mice with mCherry or NpHR in the left CeA | Unpaired t-test | P=0.0080  t=3.229 | 6-7 mice per group |
| 1N | Percent change in VMR to 60 mmHg in CYP mice with mCherry or NpHR in the right CeA before, during, and after laser stimulation | Two-way RM ANOVA with Dunnett’s posttest | Time, p=0.0002, F (1.615, 17.77) = 16.34  Virus, p=0.0008, F (1, 11) = 20.80  Time x virus, p=0.0039, F (2, 22) = 7.210  mCherry:  BL v laser, p=0.8892  BL v PL, p=0.0945  NpHR:  BL v laser, p=0.0017  BL v PL, p=0.7449 | 6-7 mice per group |
| 1O | Area under the curve for 60 mmHg pressure in CYP mice with mCherry or NpHR in the right CeA | Unpaired t-test | P=0.0007  t=4.699 | 6-7 mice per group |
| 2C | Abdominal mechanical sensitivity before and after CYP treatment | Paired t-test | P<0.0001  t=7.716 | 31 mice |
| 2D | 50% withdrawal thresholds in CYP mice with mCherry or ChR2 in the left CeA | Two-way RM ANOVA with Bonferroni posttest | Time, p=0.0103, F (1, 17) = 8.314  Virus, p=0.5377, F (1, 17) = 0.3959  Time x virus, p=0.4061, F (1, 17) = 0.7257  Post CYP v laser:  mCherry, p=0.3989  ChR2, p=0.0209 | 8-11 mice per group |
| 2E | Percent change in abdominal sensitivity in CYP mice with mCherry of ChR2 in the left CeA | Two-way RM ANOVA with Bonferroni posttest | Time, p=0.0162, F (1, 17) = 7.177  Virus, p=0.1226, F (1, 17) = 2.639  Time x virus, p=0.1226, F (1, 17) = 2.639  Post CYP v laser:  mCherry, p>0.9999  ChR2,p= 0.0083 | 8-11 mice per group |
| 2F | 50% withdrawal thresholds in CYP mice with mCherry or ChR2 in the right CeA | Two-way RM ANOVA with Bonferroni posttest | Time, p=0.1159, F (1,11) = 2.913  Virus, p=0.5393, F (1, 11) = 0.4014  Time x virus, p=0.0327, F (1, 11) = 5.963  Post CYP v laser:  mCherry, p>0.9999  ChR2, p=0.0329 | 6-7 mice per group |
| 2G | Percent change in abdominal sensitivity in CYP mice with mCherry of ChR2 in the right CeA | Two-way RM ANOVA with Bonferroni posttest | Time, p=0.0022, F (1, 11) = 15.78  Virus, p=0.0001, F (1, 11) = 32.15  Time x virus, p=0.0001, F (1, 11) = 32.15  Post CYP v laser:  mCherry, p=0.4745  ChR2, p<0.0001 | 6-7 mice per group |
| 2I | Difference score for time spent in the laser chamber in CYP mice with mCherry or ChR2 in the left or right CeA | Two-way ANOVA | Virus, p=0.4045, F (1, 29) = 0.7157  Side, p=0.6708, F (1, 29) = 0.1844  Virus x side, p=0.5057, F (1, 29) = 0.4541 | 6-9 mice per group |
| 3D | Number of spikes in left CeA neurons to increasing current steps during CGRP (500 nM) or aCSF application | Two-way ANOVA | Drug, p=0.0020, F (1, 26) = 11.77  Current injection, p<0.0001, F (1.469, 38.20) = 31.18  Drug x Current injection, p<0.0001, F (13, 338) = 6.363 | 11 mice per group |
| 3F | V_rest_ of left CeA neurons in response to aCSF or CGRP (500 nM) | Paired t-test | P<0.0001  t=9.484 | 11 mice per group |
| 3H | Number of spikes in right CeA neurons to increasing current during CGRP (500 nM) or aCSF application | Two-way ANOVA | Drug, p=0.0075, F (1, 31) = 8.183  Current injection, p<0.0001, F (1.657, 51.37) = 34.06  Drug x Current injection, p<0.0001, F (13, 403) = 4.826 | 11 mice per group |
| 3J | V_rest_ of right CeA neurons in response to aCSF or CGRP (500 nM) | Paired t-test | P=0.0367  t=2.409 | 11 mice per group |
| 3K | Percent change in VMRs to 60 mmHg distention after injection of drug into the left CeA of naïve mice | Two-way RM ANOVA with Dunnett’s posttest | Time, p=0.0021, F (3.414, 40.97) = 5.429  Drug, p=0.0012, F (1, 12) = 17.57  Time x drug, p=0.0008, F (6, 72) = 4.363  aCSF:  BL v 15, p=0.9276  BL v 30, p=0.2239  BL v 45, p=0.9954  BL v 60, p=0.0976  BL v 75, p=0.7753  BL v 90, p=0.5009  CGRP:  BL v 15, p=0.2248  BL v 30, p=0.0149  BL v 45, p<0.0001  BL v 60, p=0.2936  BL v 75, p=0.8788  BL v 90, p=0.9998 | 7 mice per group |
| 3L | Area under the curve for 60 mmHg distention after drug injection in the left CeA in naive mice | Unpaired t-test | P=0.0008  t=4.422 | 7 mice per group |
| 3M | Percent change in VMRs to 60 mmHg distention after injection of drug into the left CeA in CYP mice | Two-way RM ANOVA with Dunnett’s posttest | Time, p=0.6240, F (4.262, 144.9) = 0.6688  Drug, p<0.0001, F (3, 34) = 17.65  Time x drug, p<0.0001, F (18, 204) = 14.51  aCSF:  BL v 15, p=0.3200  BL v 30, p>0.9999  BL v 45, p=0.5062  BL v 60, p=0.7185  BL v 75, p=0.0855  BL v 90, p=0.2669  CGRP:  BL v 15, p=0.0016  BL v 30, p=0.0004  BL v 45, p<0.0001  BL v 60, p=0.0002  BL v 75, p=0.9028  BL v 90, p=0.1649  CGRP(8-37):  BL v 15, p=0.0685  BL v 30, p=0.0011  BL v 45, p=0.0051  BL v 60, p=0.0080  BL v 75, p=0.6106  BL v 90, p=0.9093  CGRP+CGRP(8-37):  BL v 15, p=0.7755  BL v 30, p=0.9807  BL v 45, p=0.9983  BL v 60, p=0.9096  BL v 75, p=0.9816  BL v 90, p=0.9717 | 9-10 mice per group |
| 3N | Area under the curve for 60 mmHg distention after drug injection in the left CeA in CYP mice | One-way ANOVA with Dunnett’s posttest | P<0.0001  F=22.16  R^2^=0.6616  aCSF v CGRP, p=0.0152  aCSF v CGRP(8-37), p<0.0001  aCSF v CGRP +CGRP(8-37), p=0.7761 | 9-10 mice per group |
| 3P | Percent change in VMRs to 60 mmHg distention after injection of drug into the right CeA of naïve mice | Two-way RM ANOVA with Dunnett’s posttest | Time, p=0.0002, F (2.216, 31.03) = 10.82  Drug, p=0.0085, F (1, 14) = 9.363  Time x drug, p<0.0001, F (6, 84) = 8.119  aCSF:  BL v 15, p=0.2393  BL v 30, p=0.8630  BL v 45, p=0.7829  BL v 60, p=0.9996  BL v 75, p>0.9999  BL v 90, p>0.9999  CGRP:  BL v 15, p=0.0442  BL v 30, p=0.1129  BL v 45, p=0.0166  BL v 60, p=0.0446  BL v 75, p=0.9995  BL v 90, p=0.8484 | 8 mice per group |
| 3Q | Area under the curve for 60 mmHg distention after drug injection in the right CeA in naive mice | Unpaired t-test | P=0.0066  t=3.151 | 8 mice per group |
| 3R | Percent change in VMRs to 60 mmHg distention after injection of drug into the right CeA in CYP mice | Two-way RM ANOVA with Dunnett’s posttest | Time, p=0.3790, F (4.259, 106.5) = 1.066  Drug, p<0.0001, F (3, 25) = 45.88  Time x drug, p<0.0001, F (18, 150) = 11.97  aCSF:  BL v 15, p=0.9600  BL v 30, p=0.6980  BL v 45, p=0.2404  BL v 60, p=0.8267  BL v 75, p=0.0170  BL v 90, p=0.0890  CGRP:  BL v 15, p=0.0440  BL v 30, p=0.0135  BL v 45, p=0.0008  BL v 60, p=0.0005  BL v 75, p=0.8480  BL v 90, p=0.2859  CGRP(8-37):  BL v 15,p=0.0244  BL v 30, p=0.0002  BL v 45, p<0.0001  BL v 60, p=0.0003  BL v 75, p=0.7076  BL v 90, p=0.9781  CGRP+CGRP(8-37):  BL v 15, p=0.2740  BL v 30, p=0.3866 BL v 45, p=0.9979  BL v 60, p=0.9983  BL v 75, p=0.9999  BL v 90, p=0.7707 | 6-8 mice per group |
| 3S | Area under the curve for 60 mmHg distention after drug injection in the right CeA in CYP mice | One-way ANOVA with Dunnett’s posttest | P<0.0001  F=53.02  R^2^=0.8642  aCSF v CGRP, p<0.0001  aCSF v CGRP(8-37), p<0.0001  aCSF v CGRP + CGRP(8-37), p=0.8942 | 6-8 mice per group |
| 4B | Percent change in VMRs to 60 mmHg after injection of drug during optogenetic stimulation in the left CeA of CYP mice | Two-way RM ANOVA with Dunnett’s posttest | Time, p<0.0001, F (1.573, 25.17) = 58.02  Drug, p=0.0003, F (2, 16) = 14.31  Time x drug, p<0.0001, F (4, 32) = 12.64  aCSF:  BL v laser, p<0.0001  BL v PL, p=0.360  CGRP(8-37):  BL v laser, p=0.5061  BL v PL, p=0.5051  AP5+NBQX:  BL v laser, p=0.0003  BL v PL, p=0.3714 | 6-7 mice per group |
| 4C | Percent change in VMRs to 60 mmHg after injection of drug during optogenetic stimulation in the right CeA of CYP mice | Two-way RM ANOVA with Dunnett’s posttest | Time, p<0.0001, F (1.167, 21.00) = 24.52  Drug, p=0.0025, F (2, 18) = 8.515  Time x drug, p=0.0008, F (4, 36) = 6.051  aCSF:  BL v laser, p=0.0286  BL v PL, p=0.3955  CGRP(8-37):  BL v laser, p=0.8161  BL v PL, p=0.3905  AP5+NBQX:  BL v laser, p=0.0077  BL v PL, p=0.5923 | 6-8 mice per group |
| 4M | Percent change in VMRs to 60 mmHg distention during a) optogenetic stimulation and b) pharmacological activation in the left CeA of CGRP heterozygous and homozygous mice treated with CYP | Two-way RM ANOVA with Bonferroni posttest | a) Time, p<0.0001, F (1.518, 22.77) = 40.12  Genotype, p=0.0709, F (1, 15) = 3.779  Time x genotype, p<0.0001, F (2, 30) = 15.06  Het v KO:  Laser, p=0.0074  PL, p=0.2824  b) Time, p<0.0001, F (2.285, 34.28) = 46.89  Genotype, p=0.2353, F (1, 15) = 1.529  Time x genotype, p=0.0994, F (6, 90) = 1.844  Het v KO:  0, p=0.1301  15, p>0.9999  30, p>0.9999  45, p>0.9999  60, p>0.9999  75, p=0.2455  90, p>0.9999 | 8-9 mice per group |
| 4N | Percent change in VMRs to 60 mmHg distention during a) optogenetic stimulation and b) pharmacological activation in the right CeA of CGRP heterozygous and homozygous mice treated with CYP | Two-way RM ANOVA with Bonferroni posttest | a) Time, p=0.0004, F (2, 26) = 10.62  Genotype, p=0.1312, F (1, 13) = 2.581  Time x genotype, p=0.0011, F (2, 26) = 8.979  Het v KO:  Laser, p=0.0004  PL, p=0.4819  b) Time, p<0.0001, F (6, 78) = 56.74  Genotype, p=0.6945, F (1, 13) = 0.1613  Time x genotype, p=0.6776, F (6, 78) = 0.6656  Het v KO:  0, p>0.9999  15, p>0.9999  30, p>0.9999  45, p>0.9999  60, p>0.9999  75, p>0.9999  90, p>0.9999 | 7-8 mice per group |
| 4P | Percent change in VMRs to 60 mmHg distention during bilateral optogenetic stimulation of PBN🡪CeA CGRP terminals in naïve and CYP-treated mice | Two-way RM ANOVA with Dunnett’s posttest | Time, p=0.1131, F (1.620, 17.82) = 2.566  Pain, p=0.22516, F (1, 11) = 1.464  Time x pain, p=0.0265, F (2, 22) = 4.304  Naïve:  BL v light, p=0.0182  BL v PL, p=0.9223  CYP:  BL v light, p=0.9792  BL v PL, p=0.8198 | 5-8 |
| 5E | Amount of CGRP measured via fluorescence intensity in the left and right CeA of CYP and saline treated mice | Two-way ANOVA with Bonferroni posttest | Side, p=0.8664, F (1, 36) = 0.3780  Treatment, p=0.137, F (1, 36) = 6.714  Side x treatment, p=0.4701, F (1, 36) = 0.5329  CYP v saline:  Left, p=0.0489  Right, p=0.3930 | 10 mice per group, 3-5 sections per mouse |
| 5N | Percent of total cells expressing *Calcrl* in the left and right CeC in CYP and saline treated animals | Two-way ANOVA | Side, p=0.5810, F (1, 16) = 0.3174  Treatment, p=0.4543, F (1, 16) = 0.5882  Side x treatment, p=0.7078, F (1, 16) = 0.1456 | 5 mice per group, 3-6 sections per animal |
| 5O | Percent of total cells expressing *Sst* in the left and right CeC in CYP and saline treated animals | Two-way ANOVA | Side, p=0.8736, F (1, 12) = 0.0264  Treatment, p=0.9322, F (1, 12) = 0.0076  Side x treatment, p=0.6502, F (1, 12) = 0.2163 | 3-5 mice per group, 3-6 sections per animal |
| 5P | Percent of total cells expressing *Prkcd* in the left and right CeC in CYP and saline treated animals | Two-way ANOVA with Bonferroni posttest | Side, p=0.0778, F (1, 16) = 3.551  Treatment, p=0.0483, F (1, 16) = 4.571  Side x treatment, p=0.3359, F (1, 16) = 0.9845  CYP v saline:  Left, p=0.8595  Right, p=0.0835 | 5 mice per group, 3-6 sections per animal |
| 5Q | Percent of total cells expressing both *Sst* and *Prkcd* in the left and right CeC of saline and CYP-treated mice | Two-way ANOVA | Side, p=0.6393, F (1, 12) = 0.2312  Treatment, p=0.4155, F (1, 12) = 0.7114  Side x treatment, p=0.4847, F (1, 12) = 0.5200 | 3-5 mice per group, 3-6 sections/animal |
| 5R | Percent of *Calcrl*-expressing cells that also express *Sst* in the right and left CeC of mice treated with CYP and saline | Two-way ANOVA | Side, p=0.1144, F (1, 13) = 0.2863  Treatment, p=0.6049, F (1, 13) = 0.2812  Side x treatment, p=0.9858, F (1, 13) = 0.0003 | 3-5 mice per group, 3-6 sections/animal |
| 5S | Percent of *Calcrl*-expressing cells that also express *Prkcd* in the right and left CeC of mice treated with CYP and saline | Two-way ANOVA with Bonferroni posttest | Side, p=0.0183, F (1, 14) = 7.125  Treatment, p=0.0015, F (1, 14) = 15.45  Side x treatment, p=0.1190, F (1, 14) = 2.758  CYP v saline:  Left, p=0.2614  Right, p=0.0029  Left v right:  CYP, p>0.9999  Saline, p=0.0117 | 5 mice per group, 3-6 sections/animal |
| S1A | Normalized VMRs of background EMG before, during, and after laser stimulation in CYP mice with mCherry or ChR2 in the left CeA | Two-way RM ANOVA | Time, p=0.5548, F (1.458, 24.79) =0.4998  Virus, p=0.1630, F (1, 17) = 2.126  Time x virus, p=0.4398, F (2, 34) = 0.8415 | 10-11 mice per group |
| S1B | Normalized VMRs of background EMG before, during, and after laser stimulation in CYP mice with mCherry or ChR2 in the right CeA | Two-way RM ANOVA | Time, p=0.2181, F (1.935, 30.96) = 1.602  Virus, p=0.3878, F (1, 16) = 0.7883  Time x virus, p=0.4498, F (2, 32) = 0.8193 | 8-10 mice per group |
| S1C | Normalized VMRs for each distention at 30 mmHg before, during, and after optogenetic stimulation in CYP mice with mCherry or ChR2 in the left CeA | Two-way RM ANOVA with Dunnett’s posttest | Time, p=0.1117, F (4.109, 78.07) =1.931  Virus, p=0.3216, F (1, 19)= 1.036  Time x virus, p<0.0001, F (8, 152) = 4.399  mCherry:  1 v 2, p=0.7939  1 v 3, p=0.8921  1 v 4, p=0.7407  1 v 5, p=0.9949  1 v 6, p>0.9999  1 v 7, p>0.9999  1 v 8, p=0.8505  1 v 9, p=0.9978  ChR2:  1 v 2, p=0.9685  1 v 3, p=0.9997  1 v 4, p=0.0398  1 v 5, p=0.0128  1 v 6, p=0.0009  1 v 7, p=0.0123  1 v 8, p=0.2902  1 v 9, p=0.6965 | 10-11 mice per group |
| S1D | Normalized VMRs for each distention at 30 mmHg before, during, and after optogenetic stimulation in CYP mice with mCherry or ChR2 in the right CeA | Two-way RM ANOVA with Dunnett’s posttest | Time, p=0.0433, F (3.589, 57.43) = 2.723  Virus, p=0.8763, f (1, 16) = 0.02503  Time x virus, p=0.0958, f (8, 128) = 1.737 | 8-10 mice per group |
| S1E | Normalized VMRs for each distention at 60 mmHg before, during, and after optogenetic stimulation in CYP mice with mCherry or ChR2 in the left CeA | Two-way RM ANOVA with Dunnett’s posttest | Time, p=0.0165, F (4.538, 86.21) = 3.061  Virus, p=0.1701, F (1, 19) = 2.033  Time x virus, p=0.0014, F (8, 152) = 3.374  mCherry:  1 v 2, p=0.9833  1 v 3, p=0.9996  1 v 4, p=0.7947  1 v 5, p=0.9255  1 v 6, p>0.9999  1 v 7, p=0.9920  1 v 8, p=0.8707  1 v 9, p=0.3440  ChR2:  1 v 2, p=0.9815  1 v 3, p=0.9923  1 v 4, p=0.0123  1 v 5, p=0.0290  1 v 6, p=0.0810  1 v 7, p=0.9389  1 v 8, p=0.7598  1 v 9, p=0.9996 | 10-11 mice per group |
| S1F | Normalized VMRs for each distention at 60 mmHg before, during, and after optogenetic stimulation in CYP mice with mCherry or ChR2 in the right CeA | Two-way RM ANOVA with Dunnett’s posttest | Time, p=0.0554, F (4.241, 67.85) = 2.396  Virus, p=0.3716, F (1, 16) = 0.8451  Time x virus, p=0.0006, F (8,128) = 3.747  mCherry:  1 v 2, p=0.9803  1 v 3, p=0.9913  1 v 4, p=0.9679  1 v 5, p>0.9999  1 v 6, p>0.9999  1 v 7, p=0.9974  1 v 8, p>0.9999  1 v 9, p=0.9184  ChR2:  1 v 2, p=0.9998  1 v 3, p=0.8029  1 v 4, p=0.0649  1 v 5, p=0.0119  1 v 6, p=0.0121  1 v 7, p>0.9999  1 v 8, p>0.9999  1 v 9, p=0.9999 | 8-10 mice per group |
| S2A | Normalized VMRs of background EMG before, during, and after laser stimulation in CYP mice with mCherry or NpHR in the left CeA | Two-way RM ANOVA | Time, p=0.4154, F (1.714, 17.14) = 0.8857  Virus, p=0.1213, F (1, 10) = 2.867  Time x virus, p=0.7146, F (2, 20) = 0.3417 | 6-7 mice per group |
| S2B | Normalized VMRs of background EMG before, during, and after laser stimulation in CYP mice with mCherry or NpHR in the right CeA | Two-way RM ANOVA | Time, p=0.6038, F (2, 20) = 0.5175  Virus, p=0.1207, F (1, 10) = 2.877  Time x virus, p=0.1485, F (2, 20) = 2.101 | 6-7 mice per group |
| S2C | Normalized VMRs for each distention at 30 mmHg before, during, and after optogenetic stimulation in CYP mice with mCherry or NpHR in the left CeA | Two-way RM ANOVA with Dunnett’s posttest | Time, p<0.0001, F (3.117, 34.29) = 9.387  Virus, p=0.1484, F (1, 11) = 2.416  Time x virus, p<0.0001, F (8, 88) = 10.49  mCherry:  1 v 2, p=0.9061  1 v 3, p=0.6000  1 v 4, p=0.9756  1 v 5, p=0.9994  1 v 6, p=0.9999  1 v 7, p=0.7363  1 v 8, p=0.9996  1 v 9, p=0.8980  NpHR:  1 v 2, p=0.3827  1 v 3, p=0.9998  1 v 4, p=0.0161  1 v 5, p=0.0158  1 v 6, p=0.0240  1 v 7, p=0.5989  1 v 8, p=0.9858  1 v 9, p=0.8877 | 6-7 mice per group |
| S2D | Normalized VMRs for each distention at 30 mmHg before, during, and after optogenetic stimulation in CYP mice with mCherry or NpHR in the right CeA | Two-way RM ANOVA with Dunnett’s posttest | Time, p=0.0533, F (3.374, 37.12) = 2.705  Virus, p=0.2648, F (1, 11) = 1.380  Time x virus, p<0.0001, F (8, 88) = 5.600  mCherry:  1 v 2, p=0.7606  1 v 3, p=0.2422  1 v 4, p=0.6577  1 v 5, p=0.9244  1 v 6, p>0.9999  1 v 7, p=7584  1 v 8, p=0.9573  1 v 9, p=0.9060  NpHR:  1 v 2, p=0.5913  1 v 3, p=0.9931  1 v 4, p=0.0185  1 v 5, p=0.0695  1 v 6, p=0.0088  1 v 7, p=0.9777  1 v 8, p=0.9534  1 v 9, p=0.6126 | 6-7 mice per group |
| S2E | Normalized VMRs for each distention at 60 mmHg before, during, and after optogenetic stimulation in CYP mice with mCherry or NpHR in the left CeA | Two-way RM ANOVA with Dunnett’s posttest | Time, p=0.0055, F (2.900, 31,90) = 5.163  Virus, p=0.1262, F (1, 11) = 2.738  Time x virus, p<0.0001, F (8, 88) = 9.118  mCherry:  1 v 2, p=0.9945  1 v 3, p=0.8628  1 v 4, p=0.6908  1 v 5, p=0.8738  1 v 6, p=0.3761  1 v 7, p=0.9264  1 v 8, p=0.7806  1 v 9, p=0.9996  NpHR:  1 v 2, p=0.9931  1 v 3, p=0.9612  1 v 4, p=0.0013  1 v 5, p=0.0260  1 v 6, p=0.0092  1 v 7, p=0.8783  1 v 8, p=0.9951  1 v 9, p=0.5416 | 6-7 mice per group |
| S2F | Normalized VMRs for each distention at 60 mmHg before, during, and after optogenetic stimulation in CYP mice with mCherry or NpHR in the right CeA | Two-way RM ANOVA with Dunnett’s posttest | Time, p=0.0008, F (3.581, 35.81) = 6.428  Virus, p=0.5642, F (1, 10) = 0.3556  Time x virus, p<0.0001, F (8, 80) = 5.263  mCherry:  1 v 2, p=0.3911  1 v 3, p=0.2249  1 v 4, p=0.9521  1 v 5, p=0.8652  1 v 6, p=0.6299  1 v 7, p=0.3329  1 v 8, p=0.4710  1 v 9, p=0.1536  NpHR:  1 v 2, p=0.3711  1 v 3, p=0.9958  1 v 4, p=0.0214  1 v 5, p=0.0271  1 v 6, p=0.0027  1 v 7, p=0.9472  1 v 8, p=0.8827  1 v 9, p>0.9999 | 6-7 mice per group |
| S3A | Percent change in VMR to 30 mmHg in CYP mice with mCherry or ChR2 in the left CeA before, during, and after laser stimulation | Two-way RM ANOVA with Dunnett’s posttest | Time, p=0.0833, F (1.877, 35.66) = 2.711  Virus, p=0.0032,  F (1, 19) = 11.38  Time x virus, p=0.0171,  F (2, 38) = 4.536  mCherry:  BL v laser, p=0.9354  BL v PL, p=0.7133  ChR2:  BL v laser, p=0.0002  BL v PL, p=0.0043 | 10-11 mice per group |
| S3B | Percent change in VMR to 30 mmHg in CYP mice with mCherry or ChR2 in the right CeA before, during, and after laser stimulation | Two-way RM ANOVA with Dunnett’s posttest | Time, p=0.0134,  F (2, 32) = 4.947  Virus, p=0.2835,  F (1, 16) = 1.232  Time x virus, p=0.1926,  F (2, 32) = 1.735 | 10-11 mice per group |
| S3C | Percent change in VMR to 30 mmHg in CYP mice with mCherry or NpHR in the left CeA before, during, and after laser stimulation | Two-way RM ANOVA with Dunnett’s posttest | Time, p<0.0001,  F (1.951, 21.46) = 59.12  Virus, p=0.0014,  F (1, 11) = 17.87  Time x virus, p<0.0001, F (2, 22) = 68.26  mCherry:  BL v laser. P=0.9757  BL v PL, p=0.6281  NpHR:  BL v laser, p=0.0001  BL v PL, p=0.0774 | 6-7mice per group |
| S3D | Percent change in VMR to 30 mmHg in CYP mice with mCherry or NpHR in the right CeA before, during, and after laser stimulation | Two-way RM ANOVA with Dunnett’s posttest | Time, p=0.0379, F (1.584,17.42) = 4.220  Virus, p=0.0596, F (1, 11) = 4.410  Time x virus, p=0.0001, F (2, 22) = 13.94  mCherry:  BL v laser, p=0.5243  BL v PL, p=0.9664  NpHR:  BL v laser, p=0.0001  BL v PL, p=0.9981 | 6-7 mice per group |
| S4A | Percent change in VMRs to 30 mmHg after injection of drug during optogenetic stimulation in the left CeA of CYP mice | Two-way RM ANOVA with Dunnett’s posttest | Time, p<0.0001, F (1.825, 28.20) = 21.41  Drug, p=0.0162, F (2, 16) = 5.399  Time x drug, p<0.0001, F (4, 32) = 11.43  aCSF:  BL v laser, p=0.0001  BL v PL, p=0.0719  CGRP(8-37):  BL v laser, p=0.7882  BL v PL, p=0.1382  AP5+NBQX:  BL v laser, p<0.0001  BL v PL, p=0.9509 | 6-7 mice per group |
| S4B | Percent change in VMRs to 30 mmHg after injection of drug during optogenetic stimulation in the right CeA of CYP mice | Two-way RM ANOVA with Dunnett’s posttest | Time, p<0.0001, F (1.355, 25.74) = 25.78  Drug, p=0.0296, F (2, 19) = 4.260  Time x drug, p=0.0002, F (4, 38) = 7.163  aCSF:  BL v laser, p=0.0020  BL v PL, p=0.6096  CGRP(8-37):  BL v laser, p=0.9551  BL v PL, p=0.6912  AP5+NBQX:  BL v laser, p=0.0322  BL v PL, p=0.8479 | 6-8 mice per group |
| S4C | Percent change in VMRs to 30 mmHg distention during a) optogenetic stimulation and b) pharmacological activation in the left CeA of CGRP heterozygous and homozygous mice treated with CYP | Two-way RM ANOVA with Bonferroni posttest | a) Time, p=0.0036, F (1.157, 17.35) = 10.45  Genotype, p=0.1270, F (1, 15) = 0.7993  Time x genotype, p=0.0003, F (2, 30) = 10.99  Het v KO  Laser, p=0.0223  PL, p>0.9999  b) Time, p<0.0001, F (2.367, 35.50) = 21.15  Genotype, p=0.8590, F (1, 15) = 0.02365  Time x genotype, p=0.9622, F (6, 90) = 0.2397  Het v KO:  0, p>0.9999  15, p>0.9999  30, p>0.9999  45, p>0.9999  60, p>0.9999  75, p>0.9999  90, p>0.9999 | 8-9 mice per group |
| S4D | Percent change in VMRs to 30 mmHg distention during a) optogenetic stimulation and b) pharmacological activation in the right CeA of CGRP heterozygous and homozygous mice treated with CYP | Two-way RM ANOVA with Bonferroni posttest | a) Time, p=0.0044, F (2, 26) = 6.718  Genotype, p=0.2477, F (1, 13) = 1.465  Time x genotype, p=0.0058, F (2, 26) = 6.319  Het v KO:  Laser, p=0.0055  PL, p=0.8418  b) Time, p<0.0001, F (6, 78) = 32.55  Genotype, p=0.1504, F (1, 13) = 1.725  Time x genotype, p=0.9629, F (6, 78) = 0.2373  Het v KO:  0, p=0.9971  15, p=0.9972  30, p>0.9999  45, p>0.9999  60, p>0.9999  75, p=0.9999  90, p=0.9945 | 7-8 mice per group |
| S5A | Percent change in VMRs to 30 mmHg distention after injection of drug into the left CeA of naïve mice | Two-way RM ANOVA with Dunnett’s posttest | Time, p=0.0007, F (3.255, 39.06) = 6.650  Drug, p=0.0503, F (1, 12) = 4.731  Time x drug, p=0.0011, F (6, 72) = 4.225  aCSF:  BL v 15, p=0.9997  BL v 30, p=0.8623  BL v 45, p>0.9999  BL v 60, p>0.9999  BL v 75, p=0.9959  BL v 90, p>0.9999  CGRP:  BL v 15, p=0.0943  BL v 30, p<0.0001  BL v 45, p<0.0001  BL v 60, p=0.0005  BL v 75, p>0.9999  BL v 90, p=0.9997 | 7 mice per group |
| S5B | Percent change in VMRs to 30 mmHg distention after injection of drug into the right CeA of naïve mice | Two-way RM ANOVA with Dunnett’s posttest | Time, p=0.0035, F (2.929, 41.01) = 5.355  Drug, p=0.0066, F (1, 14) = 10.15  Time x drug, p=0.0005, F (6, 84) = 4.566  aCSF:  BL v 15, p=0.9982  BL v 30, p=0.8354  BL v 45, p=0.2953  BL v 60, p=0.5648  BL v 75, p=0.9988  BL v 90, p=0.7415  CGRP:  BL v 15, p=0.2248  BL v 30, p=0.0193  BL v 45, p=0.1057  BL v 60, p=0.1846  BL v 75, p=0.9988  BL v 90, p>0.9999 | 8 mice per group |
| S5C | Percent change in VMRs to 30 mmHg distention after injection of drug into the left CeA in CYP mice | Two-way RM ANOVA with Dunnett’s posttest | Time, p=0.0068, F (3.880, 131.9) = 3.762  Drug, p<0.0001, F (3, 34) = 12.95  Time x drug, p<0.0001, F (18, 204) = 7.787  aCSF:  BL v 15, p=0.4836  BL v 30, p=0.5332  BL v 45, p=0.3474  BL v 60, p=0.9774  BL v 75, p=0.3924  BL v 90, p=0.1034  CGRP:  BL v 15, p=0.0048  BL v 30, p=0.2077  BL v 45, p=0.0110  BL v 60, p=0.0003  BL v 75, p=0.7363  BL v 90, p=0.5535  CGRP(8-37):  BL v 15,p=0.0700  BL v 30, p=0.0962  BL v 45, p=0.0134  BL v 60, p=0.0190  BL v 75, p=0.1890  BL v 90, p=0.0949  CGRP+CGRP(8-37):  BL v 15, p=0.9999  BL v 30, p>0.9999 BL v 45, p=0.8021  BL v 60, p=0.9877  BL v 75, p=0.6307  BL v 90, p=0.9223 | 9-10 mice per group |
| S5D | Percent change in VMRs to 30 mmHg distention after injection of drug into the right CeA in CYP mice | Two-way RM ANOVA with Dunnett’s posttest | Time, p=0.3104, F (3.666, 88.00) = 1.215  Drug, p<0.0001, F (3, 24) = 17.70  Time x drug, p<0.0001, F (18, 144) = 6.841  aCSF:  BL v 15, p>0.9999  BL v 30, p=0.9963  BL v 45, p=0.7193  BL v 60, p=0.5881  BL v 75, p=0.7100  BL v 90, p=0.9996  CGRP:  BL v 15, p=0.1429  BL v 30, p=0.1051  BL v 45, p=0.0536  BL v 60, p=0.0811  BL v 75, p=0.2305  BL v 90, p=0.9624  CGRP(8-37):  BL v 15,p=0.0038  BL v 30, p=0.0004  BL v 45, p<0.0001  BL v 60, p=0.0118  BL v 75, p>0.9999  BL v 90, p=0.9999  CGRP+CGRP(8-37):  BL v 15, p=0.0979  BL v 30, p=0.7222 BL v 45, p=0.6789  BL v 60, p=0.9823  BL v 75, p=0.9999  BL v 90, p=0.6980 | 6-8 mice per group |
| S6A | Normalized VMRs to 30 mmHg distention after infusion of CGRP into the left and right striatum | Two-way RM ANOVA | Time, p=0.4685, F (2.139, 12.83) = 0.8228  Side, p=0.7535, F (1, 6) = 0.1081  Time x side, p=0.2097, F (6, 36) = 1.489 | 4 mice per group |
| S6B | Percent change from baseline VMRs to 30 mmHg distention after infusion of CGRP into the left and right striatum | Two-way RM ANOVA | Time, p=0.3975, F (2.126, 13.30) = 1.012  Side, p=0.8762, F (1, 6) = 0.02643  Time x side, p=0.2080, F (6, 36) = 1.494 | 4 mice per group |
| S6C | Normalized VMRs to 60 mmHg distention after infusion of CGRP into the left and right striatum | Two-way RM ANOVA | Time, p=0.5683, F (2.010, 12.06) = 0.5940  Side, p=0.4206, F (1, 6) = 0.7472  Time x side, p=0.8036, F (6, 36) = 0.5007 | 4 mice per group |
| S6D | Percent change from baseline VMRs to 30 mmHg distention after infusion of CGRP into the left and right striatum | Two-way RM ANOVA | Time, p=0.6870, F (2.408, 14.45) = 0.9023  Side, p=0.8563, F (1, 6) = 0.03576  Time x side, p=6870, F (6, 36) = 0.6535 | 4 mice per group |
| S7A | Area under the curve for 30 mmHg distention during optogenetic activation of the left CeA in homozygous mice treated with CYP to compare effect of order of activation on VMRs | Unpaired t-test | P=0.7168  t=0.3804 | 4 mice per group |
| S7B | Area under the curve for 30 mmHg distention during optogenetic activation of the left CeA in heterozygous mice treated with CYP to compare effect of order of activation on VMRs | Unpaired t-test | P=0.8698  t=0.1710 | 4 mice per group |
| S7C | Area under the curve for 30 mmHg distention during optogenetic activation of the right CeA in homozygous mice treated with CYP to compare effect of order of activation on VMRs | Unpaired t-test | P=0.5468  t=0.6384 | 3-5 mice per group |
| S7D | Area under the curve for 30 mmHg distention during optogenetic activation of the right CeA in heterozygous mice treated with CYP to compare effect of order of activation on VMRs | Unpaired t-test | P=0.3171  t=1.077 | 4-5 mice per group |
| S7E | Area under the curve for 30 mmHg distention during pharmacological activation of the left CeA in homozygous mice treated with CYP to compare effect of order of activation on VMRs | Unpaired t-test | P=0.9650  t=0.0457 | 4 mice per group |
| S7F | Area under the curve for 30 mmHg distention during pharmacological activation of the left CeA in heterozygous mice treated with CYP to compare effect of order of activation on VMRs | Unpaired t-test | P=0.8906  t=0.1435 | 4 mice per group |
| S7G | Area under the curve for 30 mmHg distention during pharmacological activation of the right CeA in homozygous mice treated with CYP to compare effect of order of activation on VMRs | Unpaired t-test | P=0.2901  t=1.160 | 3-5 mice per group |
| S7H | Area under the curve for 30 mmHg distention during pharmacological activation of the right CeA in heterozygous mice treated with CYP to compare effect of order of activation on VMRs | Unpaired t-test | P=0.2730  t=1.190 | 4-5 mice per group |
| S7I | Area under the curve for 60 mmHg distention during optogenetic activation of the left CeA in homozygous mice treated with CYP to compare effect of order of activation on VMRs | Unpaired t-test | P=0.7902  t=0.2783 | 4 mice per group |
| S7J | Area under the curve for 60 mmHg distention during optogenetic activation of the left CeA in heterozygous mice treated with CYP to compare effect of order of activation on VMRs | Unpaired t-test | P=0.8289  t=0.2258 | 4 mice per group |
| S7K | Area under the curve for 60 mmHg distention during optogenetic activation of the right CeA in homozygous mice treated with CYP to compare effect of order of activation on VMRs | Unpaired t-test | P=0.1997  t=1.441 | 3-5 mice per group |
| S7L | Area under the curve for 60 mmHg distention during optogenetic activation of the right CeA in heterozygous mice treated with CYP to compare effect of order of activation on VMRs | Unpaired t-test | P=0.0647  t=2.190 | 4-5 mice per group |
| S7M | Area under the curve for 60 mmHg distention during pharmacological activation of the left CeA in homozygous mice treated with CYP to compare effect of order of activation on VMRs | Unpaired t-test | P=0.9734  t=0.0348 | 4 mice per group |
| S7N | Area under the curve for 60 mmHg distention during pharmacological activation of the left CeA in heterozygous mice treated with CYP to compare effect of order of activation on VMRs | Unpaired t-test | P=0.3134  t=1.100 | 4 mice per group |
| S7O | Area under the curve for 60 mmHg distention during pharmacological activation of right CeA in homozygous mice treated with CYP to compare effect of order of activation on VMRs | Unpaired t-test | P=0.5258  t=0.6733 | 3-5 mice per group |
| S7P | Area under the curve for 60 mmHg distention during pharmacological activation of the right CeA in heterozygous mice treated with CYP to compare effect of order of activation on VMRs | Unpaired t-test | P=0.9478  t=0.0678 | 4-5 mice per group |
| S8A-B  S8E-F | Percent of cells expressing *Calcrl* that also express *Prkcd* across the rostral-caudal axis in the left and right CeA of saline and CYP- treated mice | Mixed-effects model with Tukey’s multiple comparisons | Position, p=0.8188 F (5, 35) = 0.4382 Treatment, p=0.0009, F (1, 7) = 30.42 Side, p=0.0488, F (1, 11) = 4.907 Position x treatment, p=0.2868, F (5, 11) = 1.433 Position x side, p=0.0750, F (5, 11) = 2.755 Treatment x side, p=0.0429, F (1, 11) = 5.238 Position x treatment x side, p=0.4566, F (5, 11) = 1.009 -1.22 vs. -1.34, p=0.9936; -1.22 vs. -1.46, p>0.9999; -1.22 vs. -1.58, p=0.9997; -1.22 vs. -1.70, p=0.9660; -1.22 vs. -1.82, p=0.9993; -1.34 vs. -1.46, p=0.9714; -1.34 vs. -1.58, p=0.9481; -1.34 vs. -1.70, p=0.7173; -1.34 vs. -1.82, p=0.9479; -1.46 vs. -1.58, p>0.9999; -1.46 vs. -1.70, p=0.9884; -1.46 vs. -1.82, >0.9999; -1.58 vs. -1.70, p=0.9637; -1.58 vs. -1.82,p=0.9943; -1.70 vs. -1.82, p=0.9985 | 5 mice per group, 3-6 sections/animal |
| S8C-D  S8G-H | Percent of cells expressing *Calcrl* that also express *Sst* across the rostral-caudal axis in the left and right CeA of saline and CYP-treated mice | Mixed-effects model with Tukey’s multiple comparisons | Position, p=0.9762 F (5, 19) = 0.1540 Treatment, p=0.6989, F (1, 19) = 0.1542 Side, p=0.7641, F (1, 19) = 0.0927 Position x treatment, p=0.2655, F (5, 19) = 1.410 Position x side, p=0.0069, F (5, 19) = 4.531 Treatment x side, p=0.6187, F (1, 1) = 0.4660 Position x treatment x side, p=0.4145, F (5, 1) = 2.947 -1.22 vs. -1.34, p=0.9920; -1.22 vs. -1.46, p>0.9999; -1.22 vs. -1.58, p>0.9999; -1.22 vs. -1.70, p=0.9940; -1.22 vs. -1.82, p>0.9999; -1.34 vs. -1.46, p=0.9921; -1.34 vs. -1.58, p=0.9883; -1.34 vs. -1.70, p=0.9998; -1.34 vs. -1.82, p=0.9744; -1.46 vs. -1.58, p>0.9999; -1.46 vs. -1.70, p=0.9994; -1.46 vs. -1.82, >0.9999; -1.58 vs. -1.70, p=0.9989; -1.58 vs. -1.82, p>0.9999; -1.70 vs. -1.82, p=0.9956 | 3-5 mice per group, 3-6 sections/animal |
| S9 | Concentration of cAMP in the left and right CeA after infusion of CGRP or aCSF | Two-way ANOVA | Hemisphere, p=0.9398, F (1, 14) = 0.005911  Drug, p=0.0002, F (1, 14) = 25.25  Hemisphere x drug, p=0.7519, F (1, 14) = 0.1040 | 4-5 mice per group |

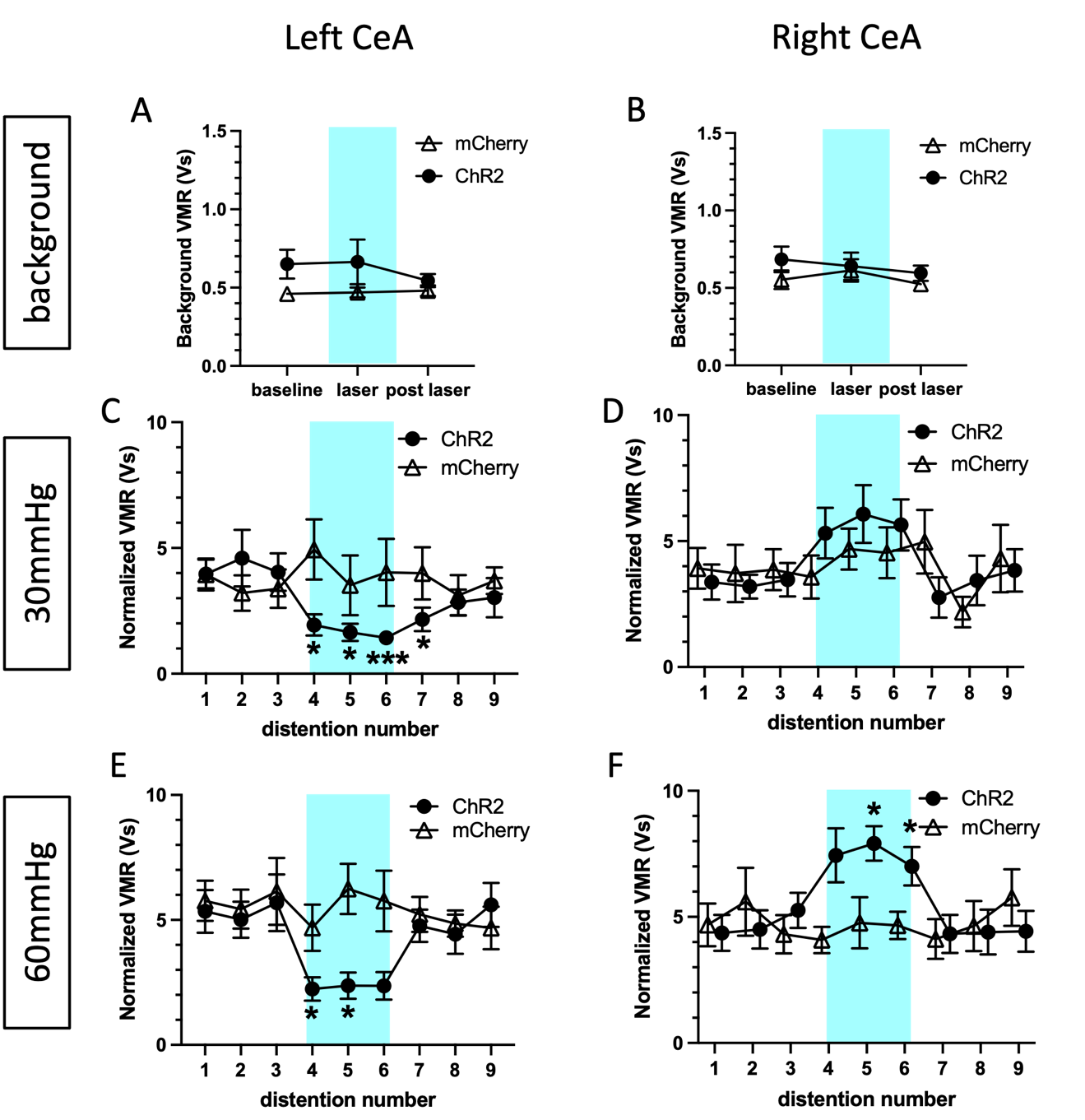

**Figure S1: Background and normalized VMRs to 30 and 60 mmHg bladder distention during optogenetic activation.** Background VMRs at baseline, during, and after optogenetic activation in the left (**A**) and right (**B**) CeA. Normalized VMRs to 30mmHg distentions in the left (**C**) and right (**D**) CeA. Normalized VMRs to 60mmHg distentions in the left (**E**) and right (**F**) CeA. All data are presented as mean +/- SEM and error bars represent SEM. *P<0.05 ***P<0.001. See **Supplementary Table 1** for further statistical information.

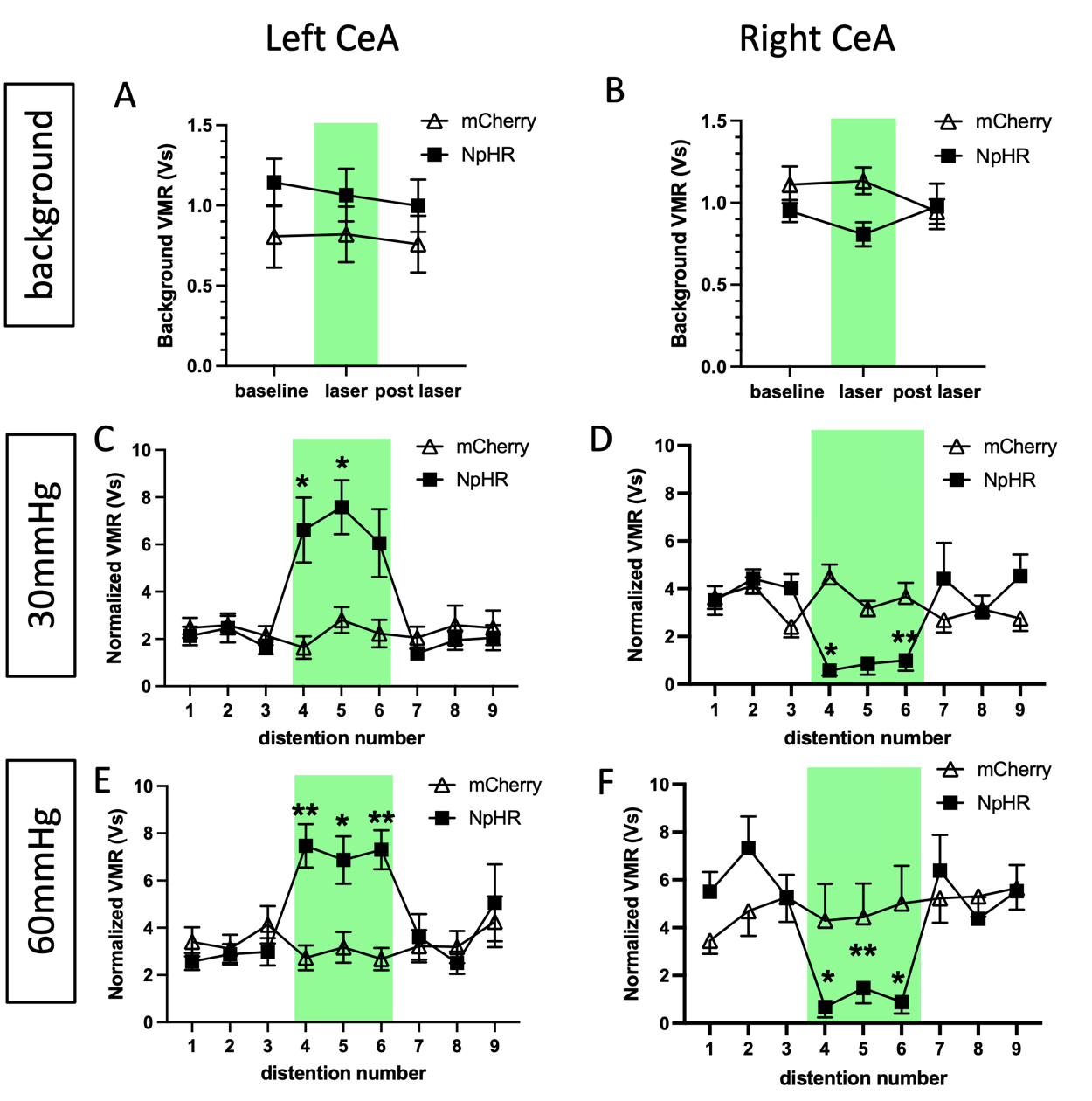

**Figure S2: Background and normalized VMRs to 30 and 60 mmHg bladder distention during optogenetic inhibition.** Background VMRs at baseline, during, and after optogenetic inhibition in the left (**A**) and right (**B**) CeA. Normalized VMRs to 30mmHg distentions in the left (**C**) and right (**D**) CeA. Normalized VMRs to 60mmHg distentions in the left (**E**) and right (**F**) CeA. All data are presented as mean +/- SEM and error bars represent SEM. *P<0.05 **P<0.01. See **Supplementary Table 1** for further statistical information.

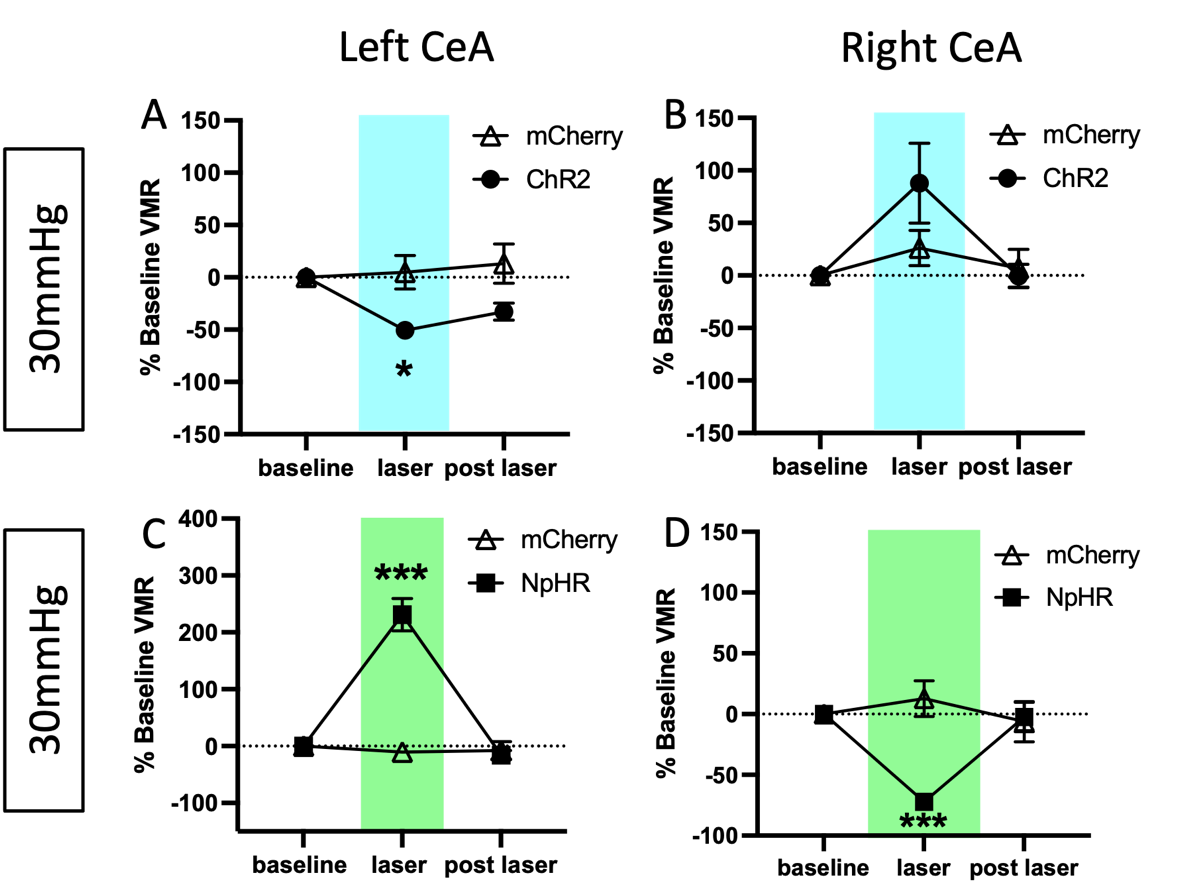

**Figure S3: Optogenetic stimulation of CGRP terminals in the left or right CeA has opposing effects on to low pressure distention.** Percent change from baseline VMRs during and after optogenetic activation of CGRP terminals in the left (**A**) and right (**B**) CeA. Percent change from baseline VMRs during and after optogenetic inhibition of CGRP terminals in the left (**C**) and right (**D**) CeA. All data are presented as mean +/- SEM and error bars represent SEM. *P<0.05, **P<0.01 ***P<0.001. See **Supplementary Table 1** for further statistical information.

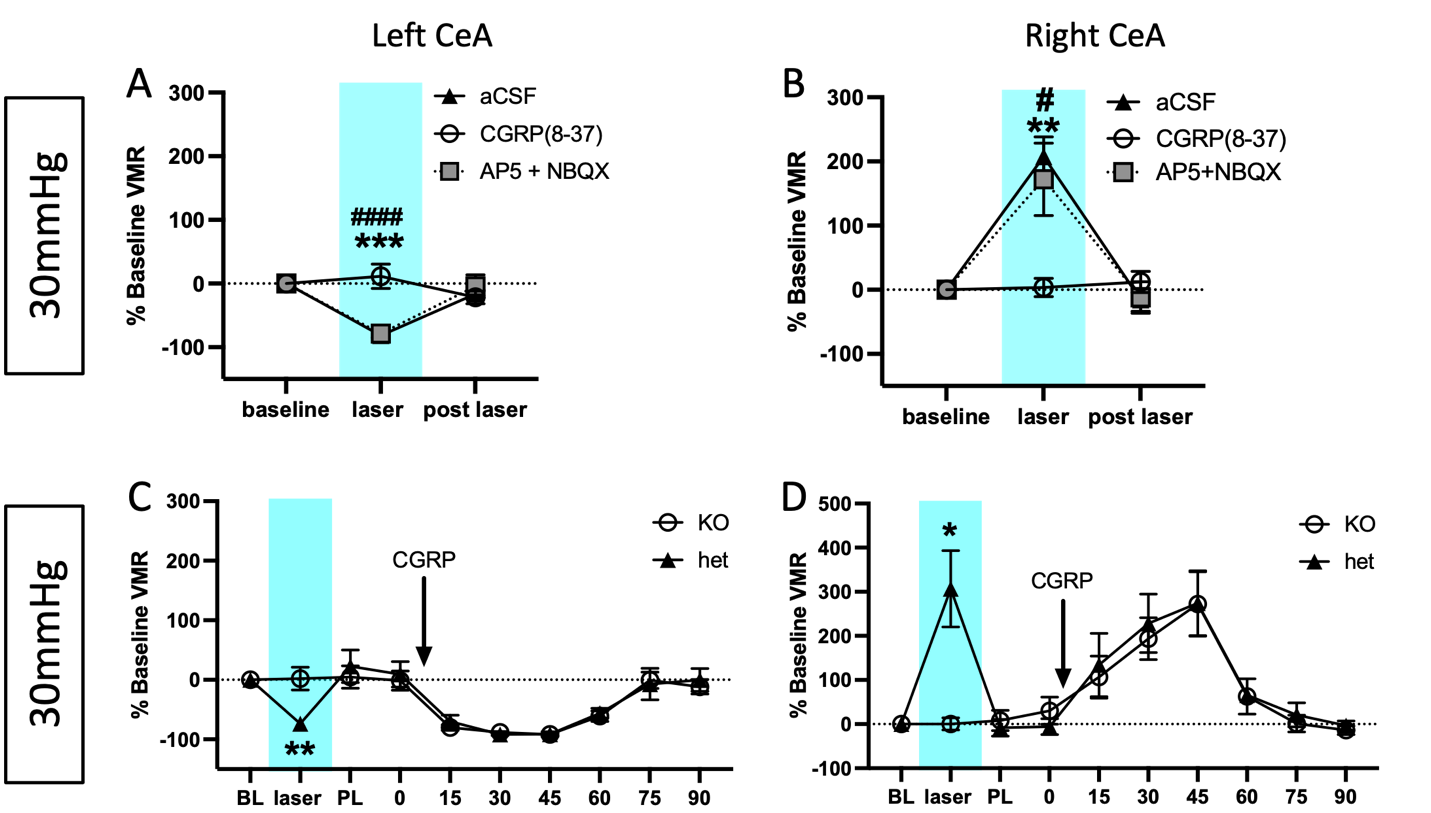

**Figure S4: Effects of optogenetic activation of CGRP terminals in the CeA during low pressure distention is due to parabrachial CGRP signaling.** Percent change from baseline VMRs to 30mmHg distention during optogenetic activation of parabrachial CGRP terminals and pharmacological inhibition of various receptor cells in the left (**A**) and right (**B**) CeA (# APv+NBQX, *aCSF). Percent change from baseline VMRs to 30mmHg distention during optogenetic and activation (left) of parabrachial Cre positive terminals and pharmacological activation (right) of CGRP receptor cells in the left (**C**) and right (**D**) CeA of *Calca-*Cre heterozygous and homozygous CYP mice. All data are presented as mean +/- SEM and error bars represent SEM. *P<0.05, **P<0.001 ***P<0.001. See **Supplementary Table 1** for further statistical information.

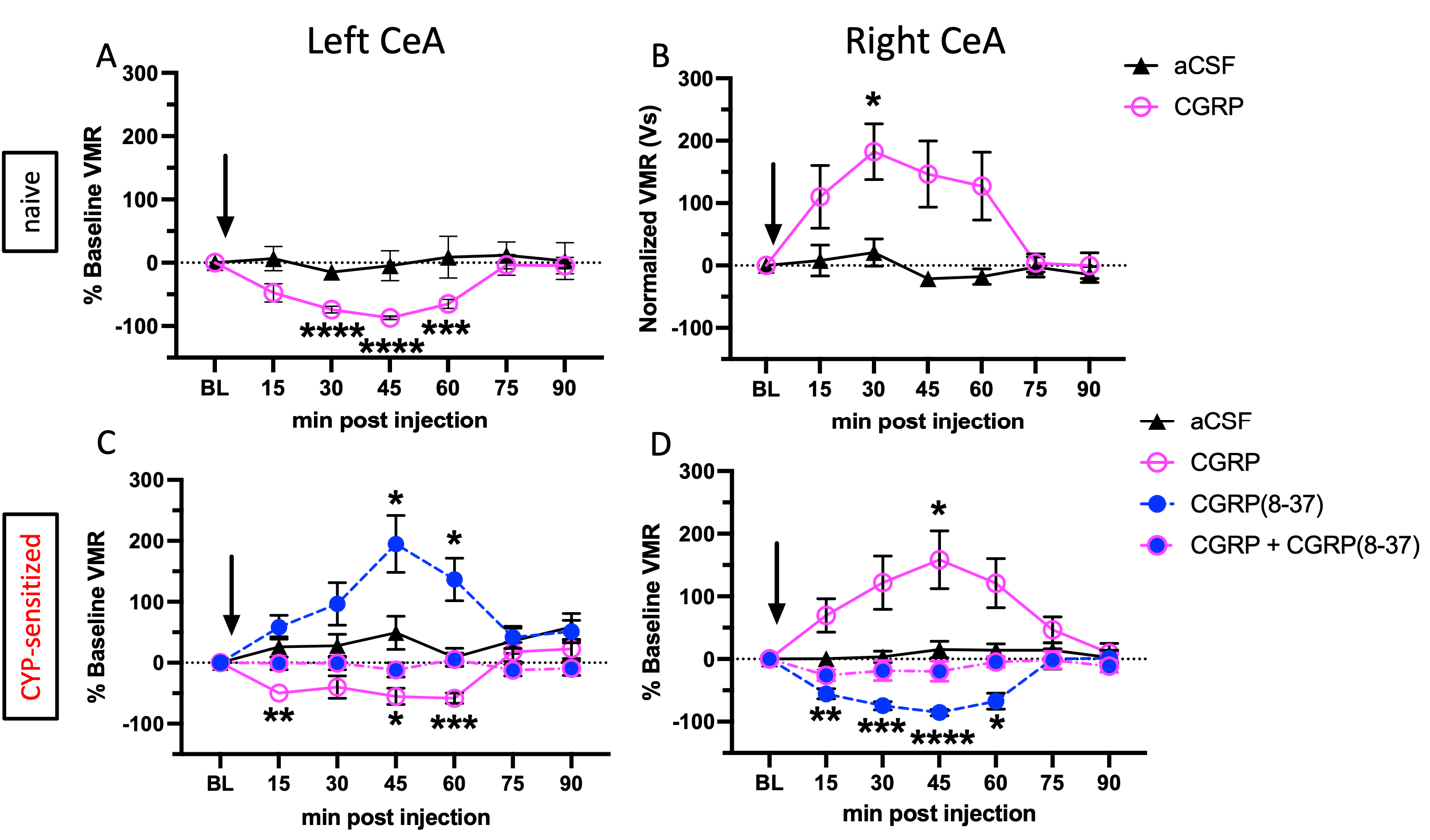

**Figure S5: CGRP pharmacology in the left and right CeA has opposing effects to low pressure distention.** Percent change from baseline VMRs to 30mmHg distention during pharmacological stimulation of CGRP receptor cells in the left (**A**) and right (**B**) CeA of naïve animals. Percent change from baseline VMRs to 30mmHg distention during pharmacological activation of CGRP receptor cells in the left (**C**) and right (**D**) CeA of CYP mice. All data are presented as mean +/- SEM and error bars represent SEM. *P<0.05, **P<0.01, ***P<0.001 ****P<0.0001. See **Supplementary Table 1** for further statistical information.

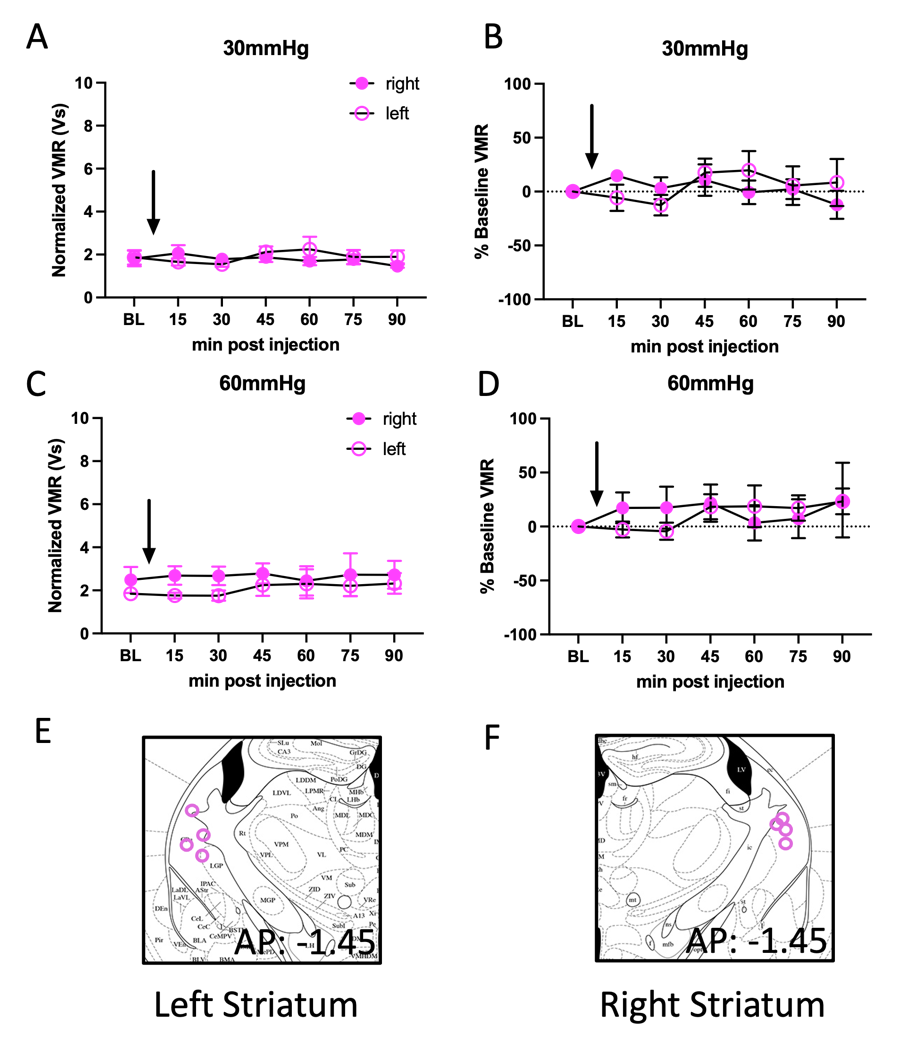

**Figure S6: CGRP pharmacology in the left and right striatum has no effect on VMRs. A**) Normalized VMRs to 30 mmHg after infusion of CGRP into the left and right striatum. **B**) Percent change from baseline VMRs to 30 mmHg distention in the left and right striatum after infusion of CGRP. **C**) Normalized VMRs to 60 mmHg distention after infusion of CGRP into the left and right striatum. **D**) Percent change from baseline VMRs to 60 mmHg distention in the left and right striatum after infusion of CGRP. **E, F**) Targeting of cannulae in the left (**E**) and right (**F**) striatum.All data are presented as mean +/- SEM and error bars represent. See **Supplementary Table 1** for further statistical information.

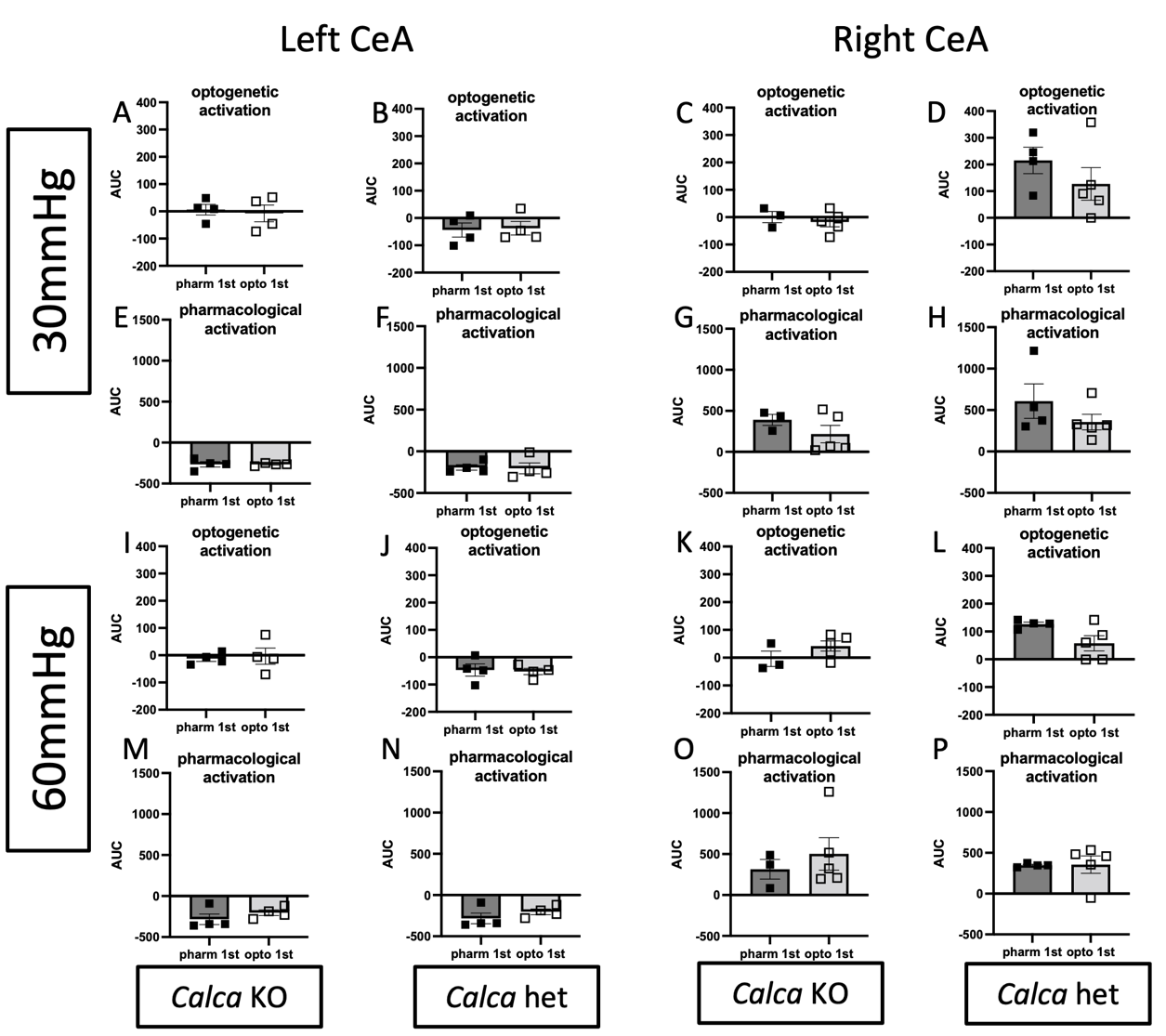

**Figure S7: Order of optogenetic or pharmacological activation has no effect on magnitude of VMR responses.** AUC for optogenetic activation in the left CeA during 30mmHg distention in *Calca* knockout (**A**) and *Calca* heterozygous (**B**) animals. AUC for optogenetic activation in the right CeA during 30mmHg distention in *Calca* knockout (**C**) and *Calca* heterozygous (**D**) animals. AUC for pharmacological activation in the left CeA during 30mmHg distention in *Calca* knockout (**E**) and *Calca* heterozygous (**F**) animals. AUC for pharmacological activation in the right CeA during 30mmHg distention in *Calca* knockout (**G**) and *Calca* heterozygous (**H**) animals. AUC for optogenetic activation in the left CeA during 60mmHg distention in *Calca* knockout (**I**) and *Calca* heterozygous (**J**) animals. AUC for optogenetic activation in the right CeA during 60mmHg distention in *Calca* knockout (**K**) and *Calca* heterozygous (**L**) animals. AUC for pharmacological activation in the left CeA during 60mmHg distention in *Calca* knockout (**M**) and *Calca* heterozygous (**N**) animals. AUC for pharmacological activation in the right CeA during 60mmHg distention in *Calca* knockout (**O**) and *Calca* heterozygous (**P**) animals. All data are presented as mean +/- SEM and error bars represent SEM. See **Supplementary Table 1** for further statistical information.

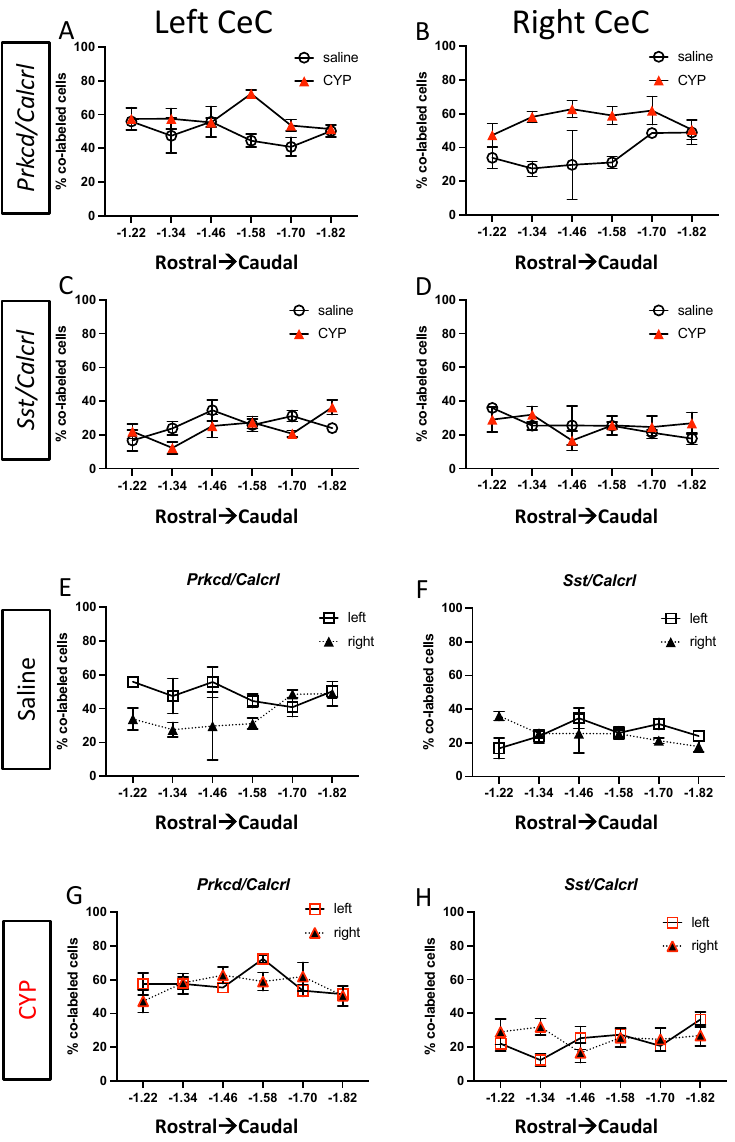

**Figure S8: Co-localization of *Sst* and *Prkcd* with *Calcrl* in the CeC based on rostral-caudal position.** Comparison of *Calcrl* co-localized with *Prkcd* between pain and non-pain animals in the left (**A**) and right (**B**) CeC. Comparison of *Calcrl* co-localized with *Sst* between pain and non-pain animals in the left (**C**) and right (**D**) CeC. Comparison of *Calcrl* co-expressed with *Prkcd* in the left and right CeC of saline (**E**) and CYP (**G**) treated mice. Comparison of *Calcrl* co-expressed with *Sst* in the left and right CeA of saline (**F**) and CYP (**H**) treated mice. All data are presented as mean +/- SEM and error bars represent SEM. See **Supplementary Table 1** for further statistical information.

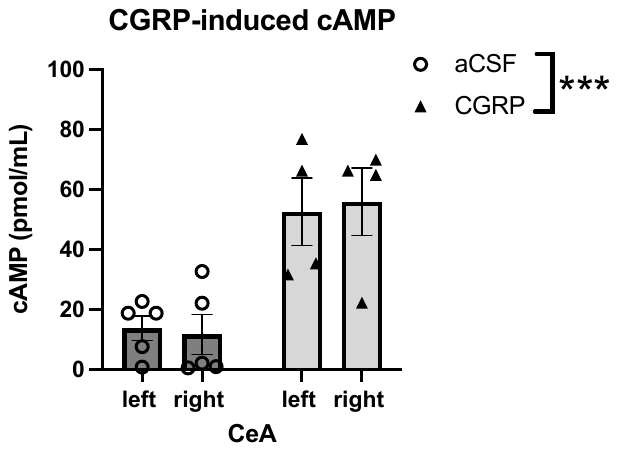

**Figure S9: Quantification of CGRP-induced cAMP in the left and right CeA.** CGRP infusion into both the left and right CeA increases cAMP compared to aCSF infusion. All data are presented as mean +/- SEM and error bars represent SEM. ***P<0.001 See **Supplementary Table 1** for further statistical information.

**References**

1. Carter ME, Soden ME, Zweifel LS, Palmiter RD (2013): Genetic identification of a neural circuit that suppresses appetite. *Nature* 503: 111–116.

2. Han S, Soleiman M, Soden M, Zweifel L, Palmiter RD (2015): Elucidating an Affective Pain Circuit that Creates a Threat Memory. *Cell* 162: 363–374.

3. Boudes M, Uvin P, Kerselaers S, Vennekens R, Voets T, De Ridder D (2011): Functional characterization of a chronic cyclophosphamide-induced overactive bladder model in mice. *Neurourol Urodyn* 30: 1659–1665.

4. Stillwell TJ, Benson RC (1988): Cyclophosphamide‐induced hemorrhagic cystitis: A review of 100 patients. *Cancer* 61: 451–457.

5. Sadler KE, McQuaid NA, Cox AC, Behun MN, Trouten AM, Kolber BJ (2017): Divergent functions of the left and right central amygdala in visceral nociception. *Pain* 158: 747–759.

6. Sadler KE, Stratton JM, Kolber BJ (2014): Urinary bladder distention evoked visceromotor responses as a model for bladder pain in mice. *Journal of Visualized Experiments*. https://doi.org/10.3791/51413

7. Ji G, Neugebauer V (2009): Hemispheric Lateralization of Pain Processing by Amygdala Neurons. *J Neurophysiol* 102: 2253–2264.

8. Tye KM, Prakash R, Kim SY, Fenno LE, Grosenick L, Zarabi H, *et al.* (2011): Amygdala circuitry mediating reversible and bidirectional control of anxiety. *Nature* 471: 358–362.

9. Chaplan SR, Bach FW, Pogrel JW, Chung JM, Yaksh TL (1994): Quantitative assessment of tactile allodynia in the rat paw. *Journal of Neuroscience Methods*, vol. 53.

10. Adke AP, Khan A, Ahn HS, Becker JJ, Wilson TD, Valdivia S, *et al.* (2021): Cell-type specificity of neuronal excitability and morphology in the central Amygdala. *eNeuro* 8: 1–28.

11. Okutsu Y, Takahashi Y, Nagase M, Shinohara K, Ikeda R, Kato F (2017): Potentiation of NMDA receptor-mediated synaptic transmission at the parabrachial-central amygdala synapses by CGRP in mice. *Mol Pain* 13: 1–11.

12. Han JS, Adwanikar H, Li Z, Ji G, Neugebauer V (2010): Facilitation of synaptic transmission and pain responses by CGRP in the amygdala of normal rats. *Mol Pain* 6. https://doi.org/10.1186/1744-8069-6-10

13. Paxinos G, Franklin KB (2019): *Paxinos and Franklin’s The Mouse Brain in Stereotaxis Coordinates*. Academic Press.
